# A simplified mesoscale 3D model for characterizing fibrinolysis under flow conditions

**DOI:** 10.1101/2023.05.09.539942

**Authors:** Remy Petkantchin, Alexandre Rousseau, Omer Eker, Karim Zouaoui Boudjeltia, Franck Raynaud, Bastien Chopard, the INSIST investigators

**Affiliations:** Scientific and Parallel Computing Group, Computer Science Department, University of Geneva, Switzerland; Complex System Modeling Group, Computer Science Department, University of Geneva, Switzerland; Laboratory of Experimental Medicine (ULB222), Faculty of Medicine, Université libre de Bruxelles, CHU de Charleroi, Belgium; Department of Neuroradiology, Hôpital Pierre Wertheimer, Hospices Civils de Lyon, Lyon, France; CREATIS Laboratory, UMR 5220, U1206, Université Lyon, INSA-Lyon, Université Claude Bernard Lyon 1, UJM-Saint Etienne, CNRS, Inserm, Lyon, France

## Abstract

One of the routine clinical treatments to eliminate ischemic stroke thrombi is injecting a biochemical product into the patient’s bloodstream, which breaks down the thrombi’s fibrin fibers: intravenous or intravascular thrombolysis. However, this procedure is not without risk for the patient; the worst circumstances can cause a brain hemorrhage or embolism that can be fatal. Improvement in patient management drastically reduced these risks, and patients who benefited from thrombolysis soon after the onset of the stroke have a significantly better 3-month prognosis, but treatment success is highly variable. The causes of this variability remain unclear, and it is likely that some fundamental aspects still require thorough investigations. For that reason, we conducted *in vitro* flow-driven fibrinolysis experiments to study pure fibrin thrombi breakdown in controlled conditions and observed that the lysis front evolved non-linearly in time. To understand these results, we developed an analytical 1D lysis model in which the thrombus is considered a porous medium. The lytic cascade is reduced to a second-order reaction involving fibrin and a surrogate pro-fibrinolytic agent. The model was able to reproduce the observed lysis evolution under the assumptions of constant fluid velocity and lysis occurring only at the front. For adding complexity, such as clot heterogeneity or complex flow conditions, we propose a 3-dimensional mesoscopic numerical model of blood flow and fibrinolysis, which validates the analytical model’s results. Such a numerical model could help us better understand the spatial evolution of the thrombi breakdown, extract the most relevant physiological parameters to lysis efficiency, and possibly explain the failure of the clinical treatment. These findings suggest that even though real-world fibrinolysis is a complex biological process, a simplified model can recover the main features of lysis evolution.

## 1 Introduction

Ischemic stroke, already reported by Hippocrates in 400 B.C.^1^, is still among the first causes of disability and death in developed countries^2^, causing around 4.5 million deaths globally per year^3^. It occurs when an aggregate of blood components clogs a brain artery, thus impeding sufficient irrigation. These thrombi, or blood clots, are mainly composed of red blood cells (RBCs) and platelets bound together by fibrin fibers, although they may contain several other biological entities. The current treatment options are either thrombolysis alone, a biochemical breakdown of the thrombus, or used with a mechanical removal of the clot (such as thrombectomy, or thrombus aspiration). The latter, whose efficacy has been demonstrated^4^ requires endovascular treatment and is indicated only in large, easily accessible arteries. The former, less invasive and applicable in first aid within a few hours after stroke symptoms onset, is done by injecting a pro-fibrinolytic macromolecule into the bloodstream. This agent, typically tissue-Plasminogen-Activator (tPA), binds to the fibrin fibers to facilitate the conversion of plasminogen to plasmin, which will then cut the fibrin strands and result in thrombus breakdown. Although the biochemistry of fibrinolysis is well understood^5^, our knowledge of many aspects of the complete process *in vivo* and *in vitro* remains sparse^6,7^. For example, the permeability of the thrombus, which is central to the transport of active macromolecules in the bloodstream, is only estimated *via* indirect measurements like perviousness^8^; the exact amount of drug that reaches the thrombus is unknown; even the dynamics of fibrin strands degradation is still actively investigated^9–14^. Even though patients who receive early intervention show a better three-month prognosis, about fifty percent fail to respond, and the reasons for this are unknown.

Hence, theoretical, experimental, and clinical research are essential to understand the underlying mechanisms of the variability of treatment outcomes and eventually improve further drug treatment development.

We present in this study a combination of experiments and modeling to investigate the dynamics of flow-driven lysis profiles of fibrin-pure clots. Experimentally, we create in a tube *in vitro* an occlusive fibrin clot in a tube, mimicking a thrombus blocking a blood vessel. We monitor the progress of the lysis and estimate the front lysis profile’s time evolution and the flow’s dynamics seeping through the clot driven by inlet and outlet pressure difference. We monitor the progress of the lysis and estimate the time evolution of the lysis profile and the dynamics of the flow seeping through the clot. The experiments show that the lysis accelerates in time. However, the flow remains constant or slightly increases during the lysis, thus suggesting that the recanalization would occur late. We translate these observations into a simple one-dimensional lysis model with constant flow and all pro-fibrinolytic agents aggregated into a single species, called anti-Fibrin Agent (anti-FA), that is blocked at the front of the clot and interacts through a second-order reaction. This model can be solved analytically and accurately predicts lysis profiles given the initial conditions.

Several numerical models have already tackled the flow-driven thrombolysis question. Most of them were inspired by Anand *et al*.’s seminal work^15^ and combined compartmental models for the temporal evolution of the proteins concentration, convection-diffusion-reaction equations to describe the transport of their free phase, and the reactions of bound proteins. Convective transport, due to blood flow, is a central issue and mainly relies on the Navier-Stokes equations, considering the clot as a homogeneous medium^16,17^ or the blood as a homogeneous visco-elastic fluid^15^, or relying on Darcy’s law with constant pressure drop^18^. Some models use either a small pressure drop ^16^, a high concentration of the profibinolytic proteins ^17,19^, or accelerated kinetics^20^. Nevertheless, they provide valuable information on simulated thrombolytic treatments, the extent of lysis, and changes in flowrate and pressure but are not directly compared to experiments realized in similar conditions.

Another approach considers multiscale fibrinolysis with microscale detailed stochastic biochemistry and lysis of a single fiber and macroscale degradation of the full clot^21^. These authors previously concluded that deterministic models of fibrinolysis might be inappropriate given the concentrations involved^22^, but the multiscale model considers only diffusional transport. A mechanistic microscale model, where the clot is modeled as red blood cells trapped in a regular assembly of elastic fibrin bonds, described the removal of clot fragments due to biochemical and mechanical degradations^23^. This fine-grained model investigates the role of flow conditions on clot fragmentation but resolves thrombolysis up to a second of physical time.

In this study, we rely on a mesoscopic approach in which the flow is computed with the open-source lattice-Boltzmann (LB) library Palabos^24^ instead of solving the Navier-Stokes equations. The fibrin clot is described as a porous material, with porosity changing locally with the progress of the lysis. Our framework can incorporate heterogeneity directly into clot porosity and composition to assess to what extent the thrombus heterogeneity can impact stroke treatment and determine whether it is possible to design novel protocols when considering thrombolysis in acute ischemic stroke.

## 2 Material and Methods

### Experimental setup

#### Clot preparation

Pasteur pipettes, with the tips cut off, were used (VWR). To form the fibrin clot at a height suitable for the experiments, we placed 900 *μ*l of glycerol (Merck) in the bottom of a tube. Then we place the pipette into the tube. The glycerol level rises in the tube to the desired level. For fibrin clot formation, we prepared solutions containing different concentrations of fibrinogen: 2.4 or 3.2 mg·ml^*−*1^ (Merck). Plasminogen (Merck) is at a final concentration of 13 *μ*g·ml^*−*1^, Owren (Stago) as a buffer. We add thrombin (Stago) at a final concentration of 2 U·ml^*−*1^ to form the clots (final volume 800 *μ*l). Then, we incubated for 15 minutes at 37°C.

#### Lysis procedure and pictures acquisition

The Pasteur pipettes, with clot inside, are then placed on a stand and a solution is added to a height of 2 cm. This consists of Owren buffer, tPA (Merck) at a final concentration of 10 ng·ml^*−*1^, FITC-labelled BSA (Abcam) at a final concentration of 0.1 mg·ml^*−*1^, plasminogen (Merck) at a final concentration of 13 *μ*g·ml^*−*1^. Lysis is induced by adding medium every 10 minutes to the mark which is 2 cm from the level of the previously formed clot. We record a photo in daylight and a photo under a UV lamp to visualize the migration of the labelled BSA.

#### Procedure to analyze the t-PA penetration in the clot

A variant of the experiment without plasminogen was performed to measure the migration of tPA. Alone the tPA is added only in the supernatant. The clot is collected after a migration time of 30, 60, and 120 minutes. The clot is then cut into four equal parts. Plasminogen is added and the whole is incubated for 16 hours. The lysate is then collected for tPA assay by Elisa (Invitrogen). In another experiment, the clot is allowed to lysate over the first quarter of its length. The clot is then removed and cut into four equal parts, left to incubate for 16 hours and the lysate is recovered for tPA determination by Elisa (Invitrogen).

#### Flow and lysis front position measurements

To monitor the evolution of the lysis, pictures of the tubes were taken every ten minutes, with the initial position of the clot marked on the tube. Each image was manually analyzed using the software ImageJ^25^, and the progression of the lysis was evaluated by measuring the distance between the initial clot position and the interface between the clot and the solution. When the interface was not horizontal, especially at the end of the lysis, the front position was measured at the middle of the interface. To evaluate the flow, we collected the liquid during successive 10-minute intervals and measured the corresponding volume that passed through the clot.

### Minimal one-dimensional model of fibrinolysis

Typical models of fibrinolysis consist of ten or more equations to account for the different reactions between species, free or bound, involved in the fibrinolytic cascade^17,18,20,21^. In^17^, the authors have shown that the number of equations necessary to describe the lysis of fibrin pure clot can be reduced to only 6. Hence, we wonder whether it is possible to lower this number even more, while preserving the main features of the lysis. To do so, we propose a one-dimensional minimal model of lysis that relies on few assumptions and that can be solved analytically. This model aims not to replace previous detailed models of fibrinolysis but rather to identify a set of minimal rules sufficient to reproduce the time evolution of the experimentally observed lysis profiles. We consider that the system is made of only two species. The first one is the fibrin (*F*), and the second, the anti-FA 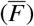, a fictive aggregate that accounts for the activation, inhibition, and binding of the different components. We further assume that the fluid velocity *u*_f_ is constant, and the anti-FA, which does not diffuse, remains blocked at the front of the clot to mimick the binding of profibrinolytic species, until the fibrin concentration decreases below a certain threshold *F*^*∗*^. We finally assume that all the profibrinolytic processes, such as activation of plasminogen into plasmin, can be included into a single reaction rate *k*_1_. The equations for the reaction and the transport of our two species finally read as:

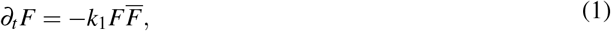

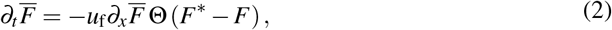

where Θ(*x*) is the Heaviside step function, with Θ(*x*) = 1 if *x >* 0 and Θ(*x*) = 0 otherwise. We write *F* and 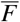 for *F*(*x, t*) and 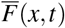 respectively for readability, and *∂*_*i*_ denotes 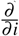. To build the model upon the observation that tPA does not penetrate deep into the bulk of the clot, we limit the penetration of anti-FA to a thickness Δ. Thus, we partition the thrombus of initial length *L* and fibrin concentration *F*_0_, into *n* slices 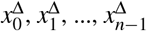 of thickness Δ. We choose the flow to be directed from the first slice 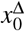 to the last 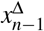 and impose a constant concentration 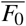 at the inlet. Under these considerations, the concentration 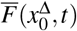 of anti-FA that accumulates at the first slice 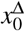 evolves as:

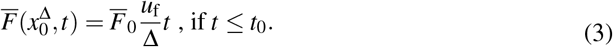

After replacing this term in eq (1) and integrating, we get:

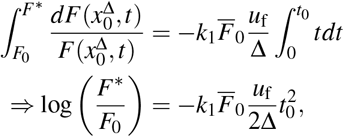

where *t*_0_ is the time to lyse the first slice, and satisfies 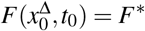. This yields:

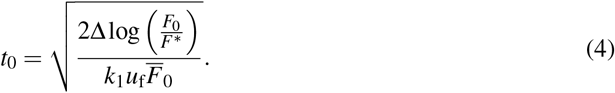

Recursively (see Section B),we obtain the time *t*_*n*_ to lyse the full clot of length *L* = *n*Δ:

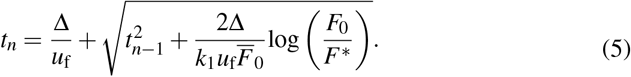

If Δ is small enough compared to the length of the clot *L*, this result is independent of Δ (see Supplementary Figure S2).

### Mesoscopic three-dimensional model of fibrinolysis

We have demonstrated that considering only one profibrinolytic species blocked at the front of the clot is sufficient to recover non-trivial lysis profiles. We now incorporate those findings in a more elaborated three-dimensional model that accounts for the fluid simulation, the thrombus as a porous medium, and the lysis. These elements are summarized in Figure 1.

**Figure 1.**
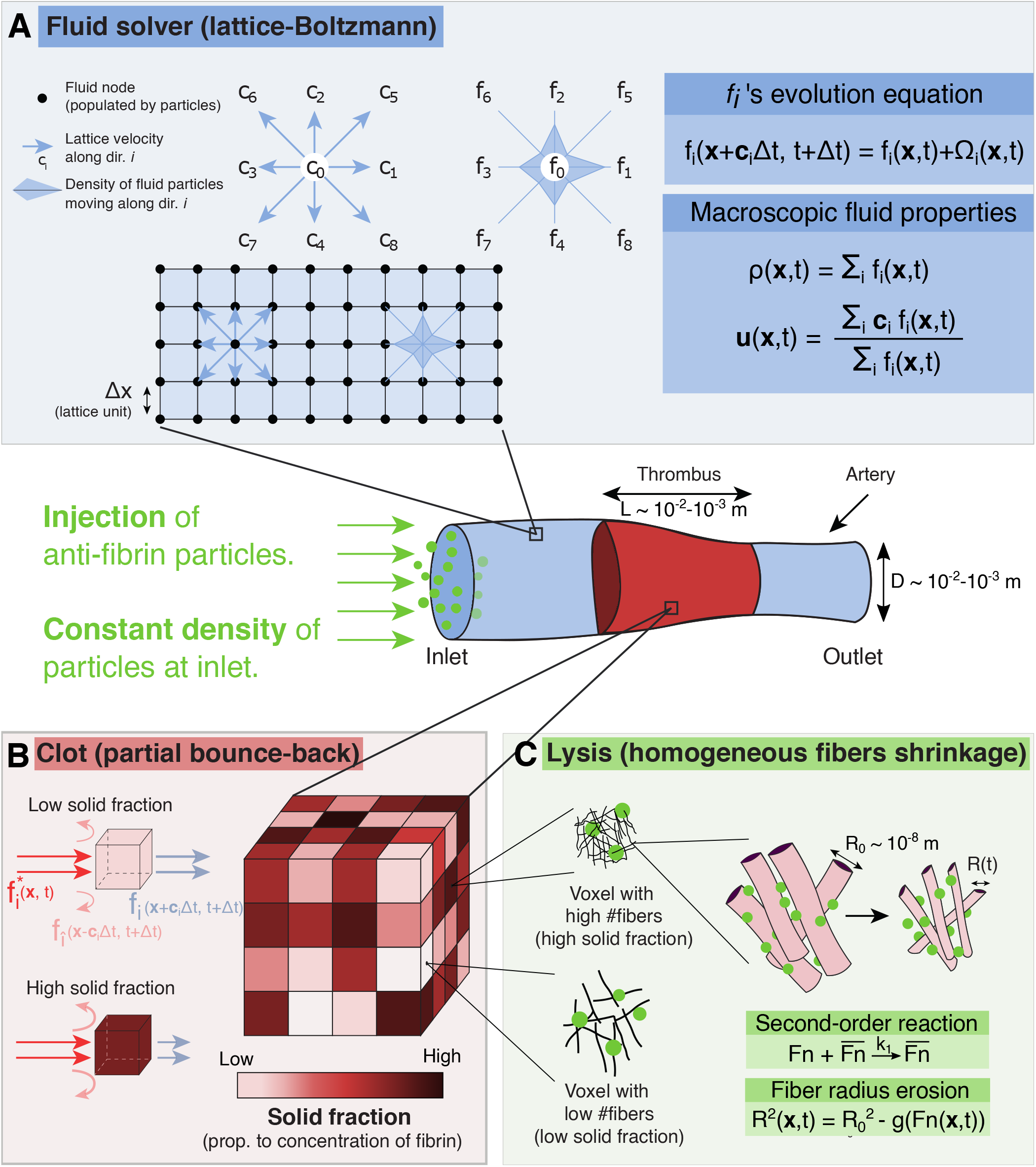
Elements of the fibrinolysis model. **A**: Fluid flow is computed in a simulated tube or vessel using the LB method. The flow is driven by a local pressure drop imposed at the left and right boundaries of the system. A clot is placed, simulated as a porous medium, using Walsh’s PBB method^26^. **B**: The blocking fraction of each clot voxel can be specified depending on the fiber volume fraction. Meso-particles of pro-fibrinolytic agents, aggregated into anti-FA, are continuously injected and will shrink the fibers’ radius upon interaction with clot voxels^18^. **C**: The lysis reaction is of second-order and relates the mean fiber radius, as well as the solid fraction in a voxel, to the fibrin concentration inside it.

#### Fluid computation

In order to simulate the fluid flow, we rely on the so-called lattice-Boltzmann (LB) method, which allows the simulation of fluids without explicitly solving the Navier-Stokes (NS) equations. The LB method can be seen as a latticed discretization of the kinetic theory, where the probability density function is replaced by fictitious particles colliding and streaming on the lattice. Inspired by Lattice Gas Automata^27^, the LB method is an efficient alternative to the NS equations^28–32^, that has been widely used to address biomedical problems^33–35^. Briefly, it solves the Boltzmann equation, which describes the evolution of so-called “fluid” particles in a regularly discretized lattice. The spatial discretization enforces a set of *q* discrete velocities *c*_*i*_, *i* = 0, 1, …, *q−*1 for the fluid particle densities *f*_*i*_, also called populations. For example, *f*_1_ is the density of fluid particles moving with velocity *c*_1_. Figure 1A depicts a 2D domain denoted D2Q9 (2 spatial dimensions and nine discrete velocities associated with nine directions). Macroscopic variables such as density *ρ* and momentum *ρ***u** are recovered by computing the moments of *f*_*i*_:

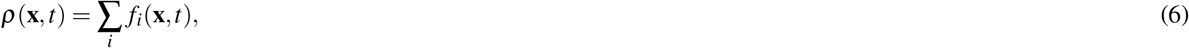

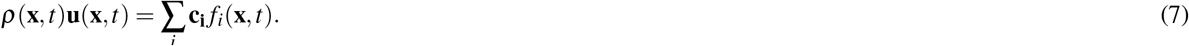

The temporal evolution of the populations *f*_*i*_(**x**, *t*) is a two steps process. First, the collision step computes at each lattice position **x** the resulting scattered populations 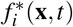. The streaming step propagates the populations 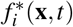 to the neighboring nodes. More formally, the scattered populations are given by:

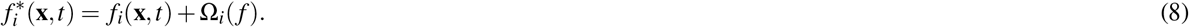

where Ω_*i*_(*f*) is the collision operator. In this paper, we consider a relaxation-type collision process between populations, the Bhatnagar-Gross-Krook (BGK) collision operator^36^:

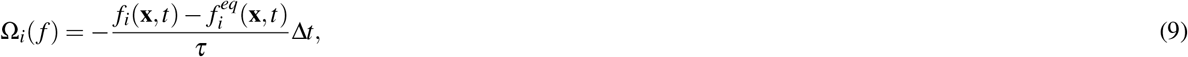

where Δ*t* is the temporal discretization, and *τ* is the model relaxation time associated with the fluid kinematic viscosity. The equilibrium populations 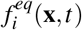 are expressed as follows (see for instance^37^):

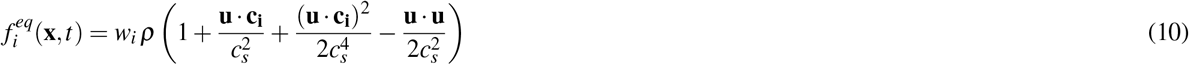

where *w*_*i*_ are weighting factors that depend on the dimension and the number of discretized velocities. *c*_*s*_ is the speed of sound in the lattice, defined by the lattice topology (see for instance^31^). Finally, the streaming step propagates the scattered populations:

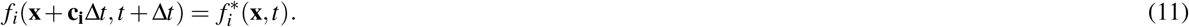

#### Clot description

Fibrinogen is a soluble blood protein made up of three pairs of polypeptide chain, and converted into fibrin by the cleavage of the fibrinopeptide by thrombin. Due to a change in solubility, fibrin monomers polymerize into protofibrils that aggregate into fibers that merge to form a branched network of fibers. The resulting fibrin fibers’ diameter range from 10^*−*8^ to 10^*−*7^ m, with pores’ diameter from 10^*−*7^ to 10^*−*5^ m^18^, and the thrombi are typically millimetric to centimetric^38^. Simulating the lysis of such an intermingled mesh at the fiber scale seems unrealistic and computationally expensive, considering the size and time scales involved. We aim to simulate the lysis of centimetric clots over hundreds of minutes while state-of-the-art detailed molecular models of fibers operate at nanometer and microsecond^39^. Instead, with a coarser description, one can treat clots as fibrous porous media^20,40,41^, which can be done in the context of LB^42^ with the so-called partial bounce-back (PBB) method.

### Partial bounce-back

When the populations *f*_*i*_ hit an obstacle during the collision step, their velocities get reversed after the streaming. This procedure leads to a zero fluid velocity relative to the boundary of the obstacle and is called solid bounce-back boundary:

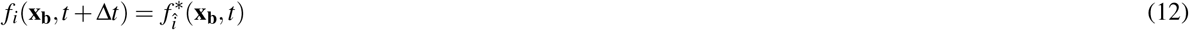

where the subscript *b* denotes a point at the boundary and *î* is the opposite lattice direction of *i*. The PBB method is an extension of the solid bounce-back boundary. The underlying idea is to assign a continuous fraction of bounce-back to each volume element, whether none, part, or all of the fluid is reflected at the boundary (Figure 1B). In LB terms, the outgoing populations in direction *i* are calculated as the weighted sum of the post-collision populations 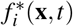 and the bounced populations 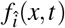^26^:

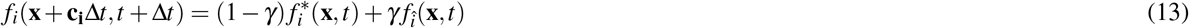

where *γ* is the fraction of bounce-back and ranges between 0 (completely fluid voxel) and 1 (completely solid voxel). Note that for *γ* = 0, we recover the usual streaming of a fluid node (Eq. (11)), while for *γ* = 1, we have the equation of the solid bounce-back boundary (Eq. (12)). Consistently, the macroscopic velocity *u*_f_ is computed as^26^:

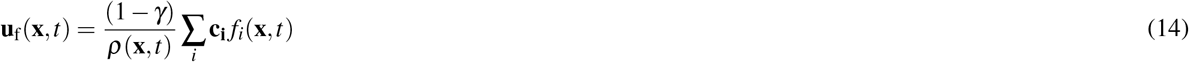

### Permeability model

The permeability is a macroscopic quantity that measures the ability of a porous medium to let a fluid flow through it. It is associated with the material’s porosity, the shape of the pores, and their local arrangements. Consequently, the permeability law of a given material is generally not known *a priori*. Instead, it has to be measured experimentally or simulated, with Darcy’s law, for instance. In^26^, the authors derived analytically the relationship between the permeability *k* and the bounce-back fraction:

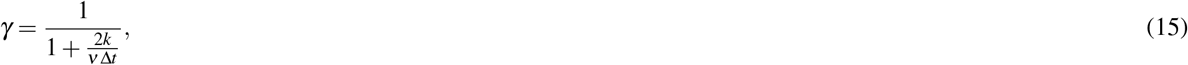

where *ν* is the kinematic fluid viscosity. However, Eq. (15) only relates the fraction of bounce-back *γ* to the permeability *k*. It implies that in order to simulate a porous material with the characteristic permeability of a thrombus, we have to incorporate in our model a permeability law that provides a value of *k*, knowing the radius of the fibers (*R*_*f*_) and the relative volume occupied by the fibers (solid fraction 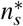). Several studies established permeability laws, either experimentally^43^ or numerically using randomly organized fibers^44^ or two-dimensional periodic square arrays of cylinders^45^. These different laws yield to similar results, but^41^ show that the permeability of *in vitro* fibrin clots is best described by Davies’ permeability law^43^:

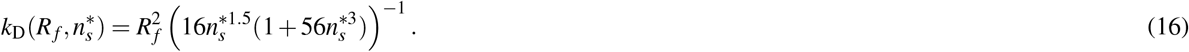

Finally, we obtain the relationship 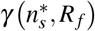 by replacing in Eq. (15) the value of *k* by Davies’ formula (16). We show (see Supplementary Figure S6) that the PBB method can be extended to accurately simulate different types of fibrous porous media over a wide range of solid fractions and reproduce the permeability law of fibrin-pure clots.

### 3D Lysis model

We previously described the clot as a collection of PBB voxels repelling the fluid with an intensity that depends on the local concentration of fibrin. To model the lysis of the clot, we use a two-species model as already proposed. Unlike the one-dimensional model, the fluid velocity is not constant but computed with the LB method that considers the clot’s permeability and the change of the solid fraction due to the decrease of fibrin concentration. The governing equations for the concentrations of the two species become:

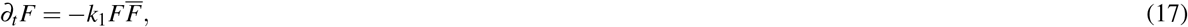

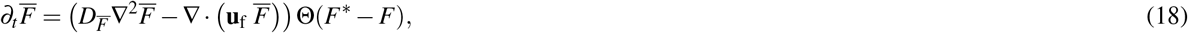

here again, we write *F* and 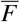 for *F*(**x**, *t*) and 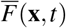 respectively for readability. For numerical reasons, we did not consider the anti-FA 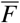 as a concentration field but treated it as an ensemble of meso-particles, each carrying a certain amount of anti-FA.

The lysis reaction occurs when meso-particles encounter fibrin voxels. The diffusion coefficient of anti-FA 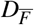 is taken as the diffusion coefficient of tPA, 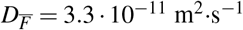^17^. The numerical resolution of equation (17) is done by finite forward differences:

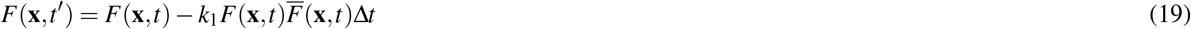

where *t*^*′*^ = *t* +Δ*t*, and the transport of anti-FA (18) is a modified version of the built-in solver of the Palabos library, see Section F.We consider that anti-FA cleaves the strands radially and uniformly^18,46,47^. As the concentration of fibrin decreases, the radius of the fibers *R*_*f*_ shrinks accordingly^18^:

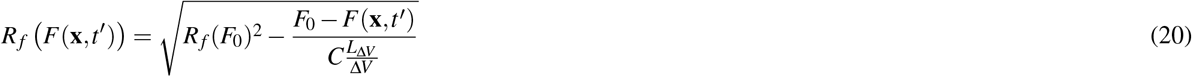

where *L*_Δ*V*_ is the total length of fibrin strands in volume Δ*V*, computed based on the fibrin concentration and fibers density, and *C* is a series of constants^18^. Then, upon reaction (19), the solid fraction 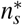 of each voxel needs to be recomputed as:

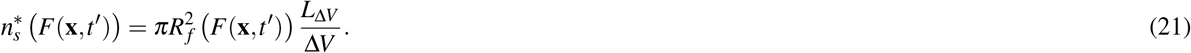

The changes in *R*_*f*_ and 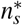 during lysis thus induce a local change in the permeability *k*, through Davies’ equation (16).

### Code availability

The numerical model is publicly available and can be downloaded from https://gitlab.com/remy.pet/insist-2020 (3D model) and https://gitlab.com/remy.pet/thrombolysis2d (2D model).

## 3 Results

We first present the experimental results, and compare them with the simplified analytical model. This also allows us to estimate a value for the model reaction parameter *k*_1_. In a second step, we show the numerical model results when varying the fibrin and anti-FA concentrations. Finally, we briefly explore the lysis of spatially heterogeneous thrombi with the numerical model.

### Penetration of tPA through fibrin pure clots

We first test the hypothesis that anti-FA remains blocked at the fibrin sites. Previous detailed molecular models already reported that due to the strong affinity of tPA with fibrin, most of the tPA is bound at the front of the clot^17,18^. However, it has not been assessed experimentally to what extent tPA remains blocked or perfuses through the clot under permeation conditions. Figure 2A schematizes the experiments where we measure the concentration of bound tPA in different regions of the clot (top most part HH, middle top part H, middle bottom part B, and bottom most part BB) after thirty minutes, sixty minutes and two hours of exposure at the clot-tPA interface. Overall, we find that tPA concentration in the HH part is significantly greater than in other parts of the clot and ten to twenty times greater than in the supernatant (Figures 2B-D). As the exposure to tPA increased, the concentration in the regions H, B, and BB increased slightly. However, it remained significantly lower than the concentration in the supernatant (Figure 2C,D). The parts of the clot in which we measure the concentration of bound tPA are considerably bigger than the regions where we assume the anti-FA to be blocked. However, we have demonstrated formally that the lysis profiles do not depend on the size of the blocking area, if it is chosen small enough (Supplementary Figure S2).For the thinner clots, the pore size can slightly hinder the transport properties of the larger molecules. We collected the solution in the vicinity of the clot front and did not measure any excess of tPA in this region (data not shown), indicating that tPA can enter the clot but is rapidly bound to fibrin. From a modeling perspective, these results support our hypothesis that anti-FA can be blocked at fibrin sites.

**Figure 2.**
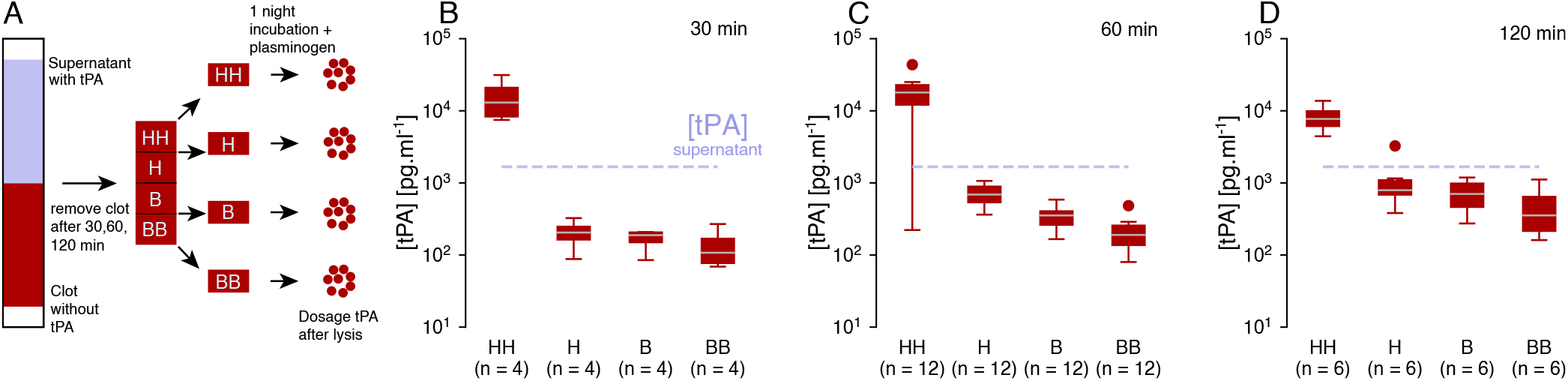
tPA does not go through the clot but remains blocked near the interface with the supernatant. **A**: Graphical sketch of the experiment. First, the supernatant containing tPA flows through the clot, then the clot is removed and cut into four pieces corresponding to the top part (HH), the top intermediate part (H), the bottom intermediate part (B), and the most bottom part (BB). Each part is lysed for one night after the addition of plasminogen, and finally, tPA concentrations are measured after centrifuging the fibrin degradation products. Concentration of tPA in different parts of the clot (HH,H,B and BB) after **B**: 30 minutes, **C**: 60 minutes and **D**: 120 minutes of perfusion. The boxplots represent the median as well as 1^st^ and 3^rd^ quartiles. The outliers are labeled as statistically non-significant, being outside of 2.689 standard deviations. The number of samples *n* varies between figures B,C and D because the corresponding experiments were independent.

### Time evolution of lysis front position and permeation

As described in the Methods, we measure the flow and the position of the clot front as the lysis proceeds. The flow is driven by pressure gradient 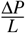, where Δ*P* is the pressure difference due to the load of the liquid column and *L* is the length of the clot. With the progress of the lysis, the clot length shortens, and liquid height increases, which should accelerate the fluid velocity, according to Darcy’s law. However, for clots prepared with 2.4 and 3.2 mg·ml^*−*1^ of fibrinogen, the flow increases moderately (Figure 3A) or stays steady during a significant part of the lysis (Figure 3C). These results are essential regarding thrombolytic treatment and recanalization. They indicate that flow is barely restored until at least half of the clot is removed. These results contrast with^17^, where simulated recanalization occurs while the average fibrin concentration in the occluded vessel is still high.

**Figure 3.**
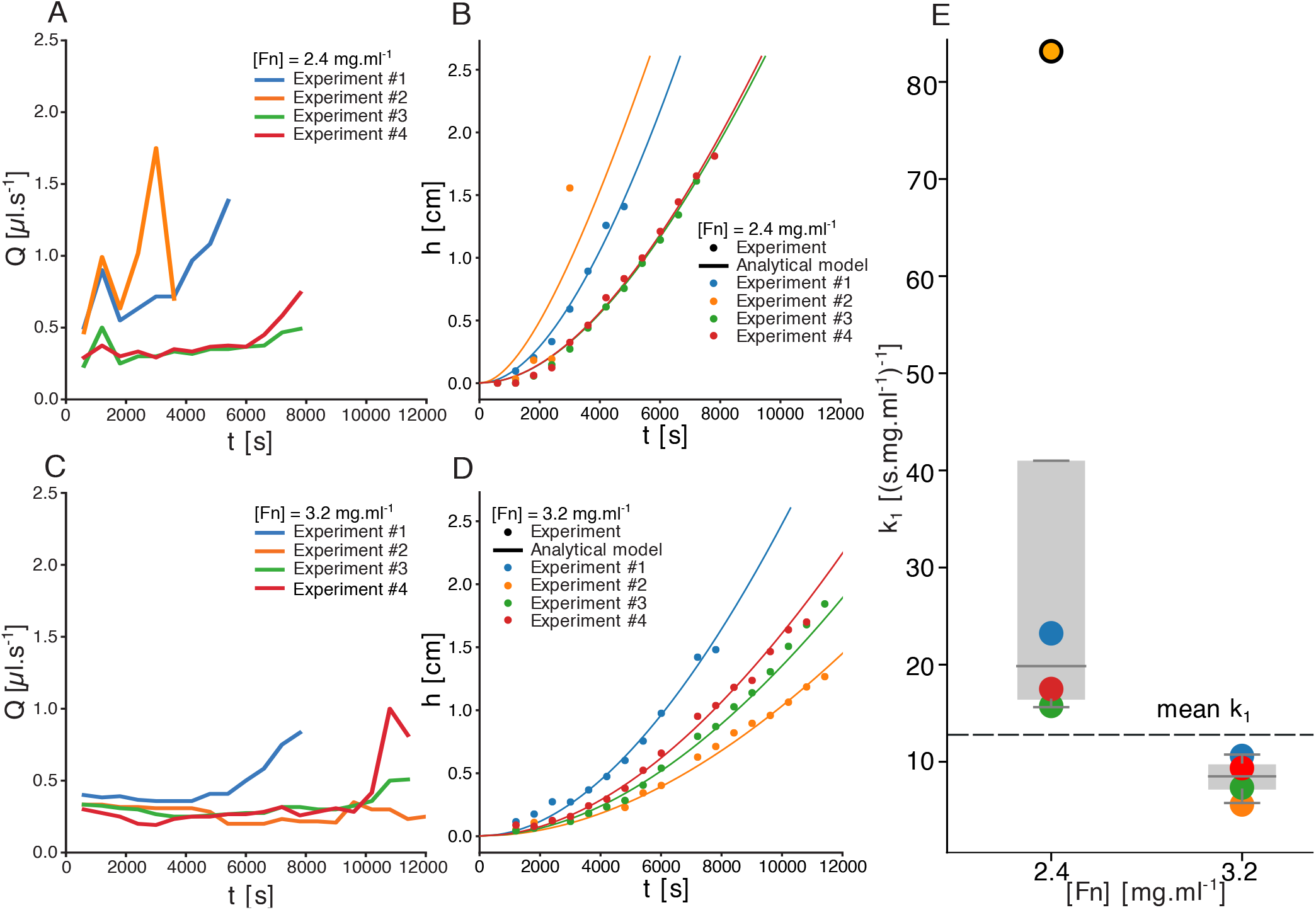
**A,C**: Flow measured in the fibrinolysis experiments of 2.4 (A) and (C) 3.2 mg·ml^*−*1^ fibrin. Experiments were repeated and numbered from 1 to 4. In most of the experiments, the flow was constant during most of the lysis and increased towards the end of lysis. **B, D**: Lysis fronts positions measured for the same lysis experiments, with 2.4 (B) and (D) 3.2 mg·ml^*−*1^ fibrin. The analytical model was fitted to the results by choosing the best value of *k*_1_. These values are reported in fig E. **E**: Best values for the analytical model’s reaction parameter *k*_1_ fitted on lysis experiments B,D. The mean value, excluding one non-statistically significant value, is 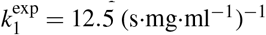. The boxplots show the 1^st^ and 3^rd^ quartiles, and the median.

Together with the flow, we also track the position of the front *h* during the lysis. Our results show that *h*(*t*) varies non-linearly with time; consequently, the lysis velocity increases as the clot is lysed. Previous numerical models found different lysis profiles, typically linear or sub-linear^16,17^. This difference can be explained by the fact that simulated bound tPA at the front of the clot tends to decrease with time, while our experimental data indicate a constant accumulation at the front (Figure 2). Despite a change in the pressure gradient, hydrodynamics effects cannot be invoked as a primary factor for lysis acceleration, considering that the flow does not change significantly during the lysis. Instead, the spatial distribution of anti-FA may be sufficient to explain the experimental lysis profiles. We measure the averaged fluid velocity for each experiment and use this value to solve iteratively equation (5). Next, we fit this solution with the experimental curves, using the corresponding values of fibrinogen and tPA concentrations (Figures 3B, D solid lines), and estimate the corresponding value of the parameter *k*_1_ (Figure 3E). For 2.4 and 3.2 mg·ml^*−*1^ of fibrinogen, we measure respectively *k*_1_ = 18.67*±*4.33 (s·mg·ml^*−*1^)^*−*1^ and *k*_1_ = 7.89*±*1.90 (s·mg ·ml^*−*1^)^*−*1^. The theoretical predictions agree with the experimental results, even though the model is oversimplified and has only one free parameter. A limitation of our model, but also present in other computational approaches, is the intrinsic variability between different experiments. Experiments performed with similar conditions result in variable outcomes for both the flow and lysis profiles, and lysis time can be up to twice longer (Figure 3). Despite identical protocol for clot preparation, the variability remains important and difficult to assess. Likely, the clot structure is involved, either local changes of the porosity or fibers radius, that may affect the flow of the degradation of the fibers. For instance, Supplementary Figure S1 shows the dispersion of the permeability among clots prepared with identical fibrin concentration. To date, analytical predictions and experimental results have not been directly compared in similar conditions, making our experimental data essential for future theoretical or numerical developments. We provide experimental lysis profiles under various experimental conditions that could be used to test and validate new models of fibrinolysis, see Supplementary Figure S3.

### Mesoscopic model of fibrinolysis

We will now show the results of the numerical model varying the fibrin and anti-FA concentrations, and briefly explore the lysis of spatially heterogeneous thrombi. Unless said otherwise, the parameters used for the simulations presented thereafter are gathered in Table 1.

**Table 1.** Typical initial conditions for the numerical and analytical models. It is specified when other values are used in the manuscript.

| Quantity | Definition | Value |
| --- | --- | --- |
| $\nu$ [m<sup>2</sup>/s] | Solution kinematic viscosity | $1 \cdot 10^{-6}$ |
| $\nabla P$ [Pa/m] | Pressure gradient across clot | $7.5 \cdot 10^3$ |
| $D$ [mm] | Tube diameter | 5.8 |
| $F$ [mg/ml] | Clot fibrin concentration | 2 – 4 |
| $F^*$ [mg/ml] | Fibrin concentration threshold | 0.2 |
| $\bar{F}$ [mg/ml] | Anti-FA concentration | $3.3 - 20 \cdot 10^{-6}$ |
| $k$ [m<sup>2</sup>] | Clot permeability | $10^{-13}$ |
| $k_1$ [(s·mg·ml<sup>-1</sup>)<sup>-1</sup>] | Estimated fibrin reaction rate | 12.5 |
| $L$ [mm] | Clot length | 5, specified else |
| $R_f$ [nm] | Fibrin fiber radius | 140 |
| $u_f$ [μm·s<sup>-1</sup>] | Mean fluid velocity (Figure 4) | 4 – 10 |
| $\Delta x$ [m] | Space discretization | $1 \cdot 10^{-4}$ |
| $\Delta t$ [s] | Time discretization | $2.5 \cdot 10^{-5}$ |

#### Homogeneous thrombi

Our LB numerical model can be used in different settings, for instance, inside patient-specific geometries (Figure 4A), or experimental tubes (Figure 4B). Our computational framework is composed of three main building blocks. The clot is described as a porous medium, the fluid computed with the LB method, and flow characteristics depend on the presence of the clot and its gradual lysis (see Methods). More importantly, our model accounts for the interplay between those blocks: with the progression of the lysis, the local concentration of fibrin decreases, which reduces the solid fraction and the permeability at a voxel level and eventually modifies globally the fluid flow as well as transport properties. Our implementation of the partial bounce-back method can reproduce predicted fluid velocity profiles (Supplementary Figure S6A) and simulate fibrous porous at any solid fraction (Supplementary Figure S6B-D).

**Figure 4.**
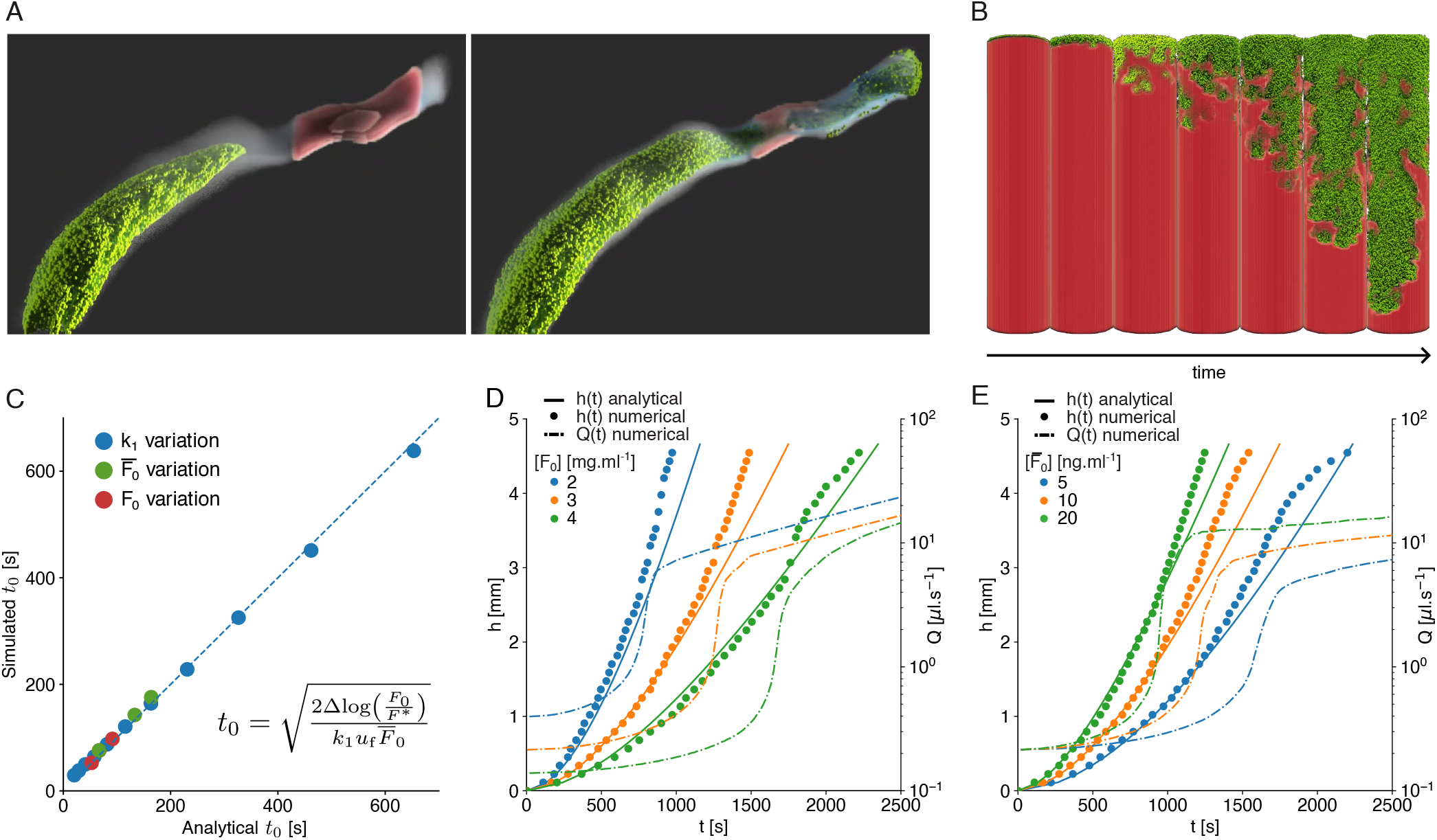
**A**: Snapshots of a fibrinolysis simulation with the 3D model in a patient-specific artery. The fibrin thrombus (red) initially blocks part of the flow (blue), which makes it hard for anti-fibrin particles (green) to reach it. When the anti-fibrin finally reaches the thrombus and does the lysis reaction, the artery can be recanalized. **B**: Snapshots of fibrinolysis with the 3D model in a cylinder in which a pressure gradient is applied. The axis shows time evolution at regular time intervals. The fibrin thrombus (red) is lysed by the anti-fibrin particles (green), following the lysis equation (17). At each time step, the view angle changes by approximately 50°. **C**: Comparison between simulated time (3D) and expected time (analytical) to lyse the first slice of the clot, obtained for different reaction parameters *k*_1_ (blue), initial concentrations of anti-FA 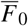 (green) and fibrin *F*_0_ (red). Here Δ = Δ*x* = 114 μm. **D, E**: Lysis front positions in time for different concentrations of fibrin and anti-FA, respectively, with the numerical 2D (dots) and the analytical (line) models. Baseline parameters are *F*_0_ = 3 mg·ml^*−*1^, 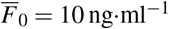, *k*_1_ = 280 (s·mg·ml^*−*1^)^*−*1^, ∇*P*_0_ = 8250 Pa·m^*−*1^, Δ*x* = 1.14 · 10^*−*4^ m, Δ*t* = 1.83 · 10^*−*4^ s.

Furthermore, we show that the minimal two-species lysis model leads to a non-linear front evolution under the assumption that one species is blocked at the clot interface. However, we must test whether such assumptions remain valid once implemented in a more complex numerical setup. For different initial concentrations of fibrinogen *F*_0_, anti-FA 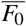 and reaction constant *k*_1_ the time to lyse the first slice of clot Δ is in agreement with the time predicted by Equation (4). The layer is considered lysed for the one-dimensional model when the fibrin concentration decreases below a threshold value *F*^*∗*^. In three dimensions, not all the voxels are lysed simultaneously due to the Poiseuille flow profile and the diffusion of the anti-FA. The layer is lysed when the average fibrin concentration in a slice goes below *F*^*∗*^, which results in small deviations from the theoretical prediction (Figure 4A). Then we simulate the complete lysis of a two-dimensional clot of width 5.8 mm and length 5 mm, with different initial concentrations of fibrin (Figure 4B) and anti-FA (Figure 4C). As observed experimentally, the lysis front velocity increases in time, and the simulated lysis profiles match the theoretical ones (see Supplementary Figure S7).Interestingly, we observe a systematic deviation from the predicted profiles in the last part of the lysis. In the last third of the simulation, the lysis is faster than expected by Equation (5). This discrepancy shows the effect of considering a constant fluid velocity. Indeed, the flow rates computed from the simulations increase initially very slowly, restoring only around 15% of the final throughput, followed by an abrupt change (Figure 4B, C dashed lines), which manifests itself by a deviation from the theoretical lysis curve. It was already reported that enhanced permeation results in faster lysis^18^. In the experiments we observe a late increase of the flow but less pronounced than in the simulations. It is worth noticing that experimental lysis profiles were not acquired until the end of the lysis. Clot front positions and flow were monitored at constant frame rate (see Methods). Whenever the measurements stopped before the lysis of the full clot, it implies that the clot was removed between two consecutive images, and likely the flow increased abruptly, as in the simulations, but could not be measured.

#### Heterogeneous thrombi

Thrombi in ischemic strokes are not homogeneous^48–51^. We take advantage of the high adaptability of the numerical mesoscopic model to explore the flow-driven lysis of heterogeneous thrombi. Clot heterogeneity can affect thrombi degradation due to a change in composition that either modifies the biochemistry of the lysis or the permeability and transport properties locally. In the present work, we focus on changes in permeability, but our LBM implementation has the versatility to consider both types of heterogeneities. It is an advantage of the mesoscopic approach that allows modifying the properties of the voxels directly, while a continuum description would rely, for example, on multiphase flows modeling^52,53^, coupling Navier-Stokes/Darcy equations with different interfaces or penalized Navier-Stokes equation^54^. The spatial heterogeneous concentration of fibrin was considered for submillimetre clots^22^ to elucidate why coarse clots lyse faster than fine clots. However, no significant differences in clot degradation rates were observed when the model was run with parameters commonly used in the literature. To some extent, the original work of Diamond *et al*.^18^, as well as subsequent studies, allow for clot heterogeneous permeability through local modification of fibrin concentration of fiber radius^19,20^ or enhanced kinetics^55^. However, this heterogeneity results from the lysis rather than the initial intrinsic features of the clots. We study two types of numerical thrombi. Type 1 thrombi are created by drawing for each site a random fibrin concentration within an uniform distribution, centered around the target concentration of 2 mg·ml^*−*1^. The standard deviation of the law comes in four variants: 25, 50, 75 and 100% of the target concentration. For instance, a thrombus generated at the target fibrin concentration of 2 mg·ml^*−*1^ and 100% dispersion, can have the concentration of each site vary between 0 and 4 mg·ml^*−*1^. Type 2 thrombi are created by randomly placing 25 medium sized-disks with a high fibrin concentration, and by filling the remaining space with a lower concentration, chosen so that the average concentration in the whole thrombus is equal to the target concentration of 2 mg·ml^*−*1^. The disks can overlap each other or can be cut if they exceed the domain’s boundary. This means that there are thrombi that have disks occupying more space than others, which implies that they have a lower outer-disk fibrin concentration to keep the same average target concentration. We imposed three intra-disk concentrations : 2.5, 3 and 3.5 mg·ml^*−*1^, corresponding to an increase of respectively 25, 50 and 75% of the target concentration.

For type 1 clots, we generated and lysed 10 thrombi for each fibrin dispersion percentage. Since the lysis front is no longer a good metric for heterogeneous thrombi, due to the heterogeneity of each slice of thrombus, we measure lysis progression by computing the thrombus remaining mass percentage and recanalization, see Figure 5A and B. We can see that the more heterogeneous the thrombi, the earlier the recanalization starts (*∼*10% faster in average) and the more variability can be observed in the recanalization profile. One of the simulation at 100% standard deviation recanalizes only one tenth of the final flow of the homogeneous simulation after 500 s. Lyses that have a lower final outflow typically have clot remnants on the walls or in initially high concentration spots. The times to lyse 25, 50 and 75% of the clot, 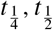 and 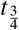 respectively, are reported in Figure 5C. Thrombi that are more heterogeneous lyse on average faster, and show more variability in lysis times. Finally, the lysis velocities standard deviation tends to increase in time. This can be seen from the boxplot Figure 5C, where we can see the quartiles of the boxes vary in time.

**Figure 5.**
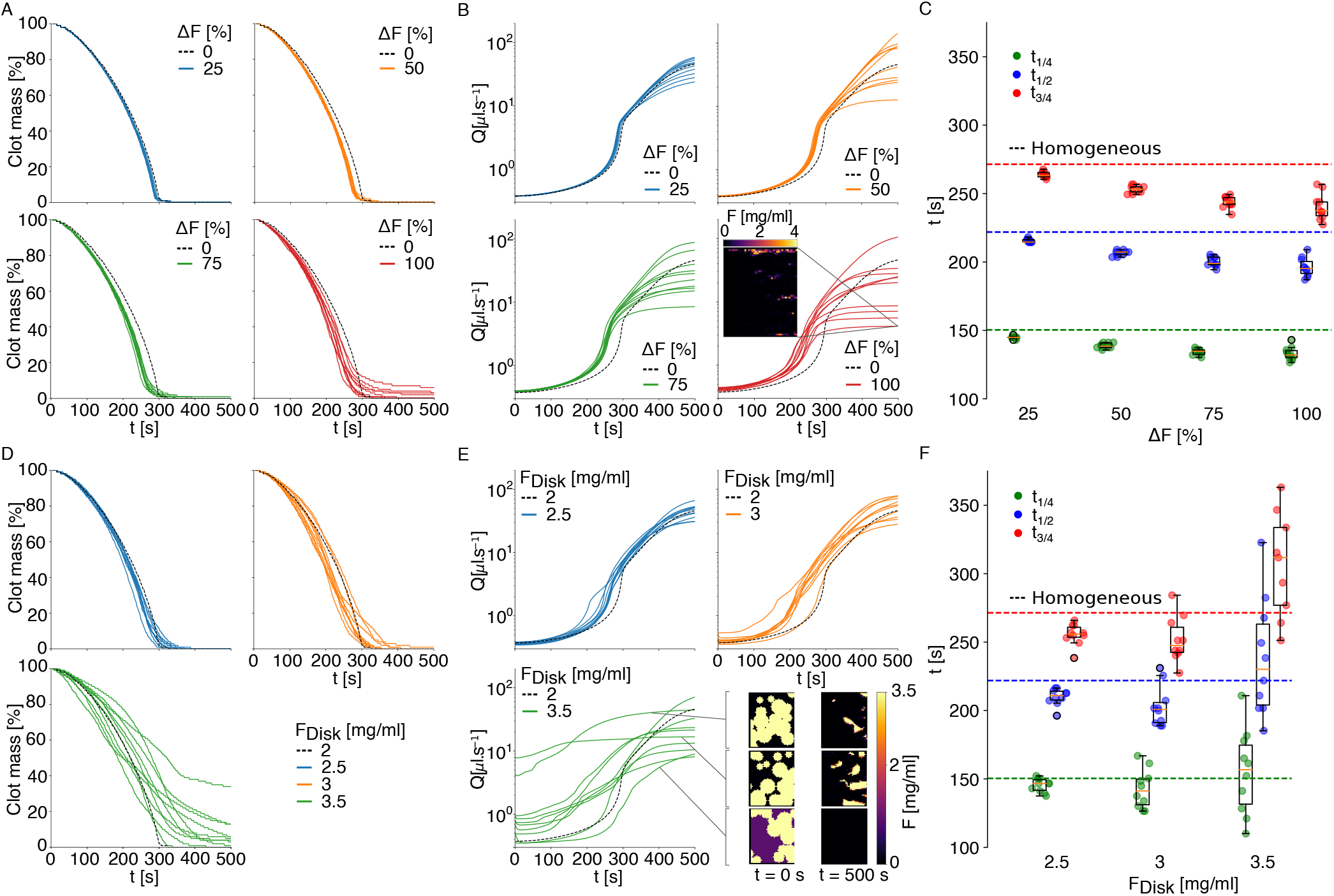
10 type 1 (A-C) uniformly heterogeneous 2D thrombi for each fibrin standard deviation, and 10 type 2 (D-F) heterogeneous 2D thrombi, for each intra-disk concentrations of 2.5, 3 and 3.5 mg·ml^*−*1^, of length 5 mm and width 5.8 mm were lysed, with ∇*P* = 8000 Pa·m^*−*1^ and *k*_1_ = 1400 (s·mg·ml^*−*1^)^*−*1^. The mean fibrin concentration for each clot is *F*_*M*_ = 2 mg·ml^*−*1^, and the initial anti-FA 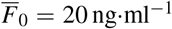. **A**,**D**: The remaining clot mass is plotted in time. **B**,**E**: The average throughput inside the clot. **C**,**F**: The times to reach 25, 50 and 75% of the lysis are plotted. The more heterogeneous the clots are, the higher is the variablity on the lysis times. One simulation with *F*_*Disk*_ = 3.5 mg·ml^*−*1^ did not reach the 75% lysis threshold during the simulation time, and was excluded from this boxplot.

For type 2 thrombi, we lysed three sets of 10 thrombi for each of the three intra-disk fibrin concentrations: 2.5, 3, and 3.5 mg·ml^*−*1^. As for the previous type of heterogeneity, we see that the more heterogeneous the thrombus, the more variability there is in the lysis curves, see Figure 5D and F, and the less abrupt their recanalization occurs, see Figure 5E. These variations are more pronounced than for type 1 clots, with fluctuation of *±*35% on half-lysis time for the most heterogeneous case. Also, the lower the outer-disk concentration, the higher the average flow inside the thrombus before the lysis, but the harder it will be to accumulate anti-FA at the thrombus disks to lyse them. This is because the higher permeability outer-disks regions act as preferred lesser resistance channels for the flow. This can produce lysis curves that reach a plateau, with a significant fraction of clot mass remaining. More variability on the lysis times and fluxes can be observed with type 2 clots than type 1 clots, see Figure 5F.

## 4 Discussion

We presented original *in vitro* fibrinolysis experiments of homogeneous fibrin-pure thrombi perfused continuously with tPA, and an analytical formula that could predict the lysis front evolution, provided the effective reaction rate *k*_1_ is well calibrated. These revealed three key points: (1) tPA does not penetrate in the bulk of the fibrin-pure thrombi, (2) the lysis accelerates, and (3) flow is restored suddenly only at the end of the lysis. We also presented an original numerical framework that uses the LB and the PBB methods to predict more specifically the evolution of clot length, permeability, and hydrodynamic properties during the fibrinolysis. Due to a strong affinity with fibrin, our experimental results show a tPA concentration profile characterized by an excess at the front of the clot and a lack, compared to the supernatant concentration, at the back. This localization of tPA leads to gradual lysis from the front to the back. Interestingly, the experimental lysis patterns differed from the simulated ones observed in ^16,19^, where the permeation of proteins through the clot and decreased bulk resistance started before front lysis. This discrepancy could be explained by a more considerable pressure drop in the simulations compared to our experimental conditions. It would be interesting to consider novel experiments with higher pressure gradients to assess whether there is a transition between sharp tPA localization with front-to-back lysis and clot degradation occurring both at the surface and in bulk.

The lysis front evolution matched the analytical model while the flow throughout lysis stayed of the same order of magnitude as the initial flow. Our results reveal that the fibrin concentration and permeability of the thrombus, dictating the transport of pro-fibrinolytic macromolecules to the fibers, is of utmost importance for fibrinolysis success.

The spatial thrombus heterogeneity alone showed a significant influence on lysis time and recanalization, with up to *∼*60% variation in half-lysis time for different instances of type 2 clots, containing high concentration disks of variable size. We could imagine that other internal arrangements of thrombi would make an even greater difference, and it would be interesting to study which topology lyses the fastest. To study more realistic thrombi, it is possible with the present model to generate and lyse clots reconstructed from real clot images. Indeed, thrombi analyzed after a thrombectomy show a very large heterogeneity in the spatial distribution of the blood’s figurative elements but also in the structure of the fibrin network^56^. All these elements are not easy to take into account in the design of an experiment. In addition to this spatial heterogeneity, there is the influence of cell surfaces in the control of fibrinolysis. Whether it is the receptors specific to the factors involved in fibrinolysis^57^ or annexin A2^58^. What further complicates the modeling is that one cell type can have completely opposite effects on fibrinolysis. We have recently shown that platelets can have a profibrinolytic or antifibrinolytic action depending on the clinical situation of the patients^6^. Numerical experimentation will be a complementary tool to address these issues. A showcase example can be found in the Supplementary Figure S8, where we lysed 2D slices of patient retrieved thrombi (sliced and analyzed by Staessens *et al*.^56^). 3D reconstructed thrombi could also be lysed, provided 3D data is available. We chose in the present work to treat heterogeneity as a change in fibrin concentration, but other options could be to change locally the reaction rate *k*_1_, or the permeability (fiber radius or permeability law), which is feasible with our numerical model.

The numerical model is purposely streamlined, keeping only the essential features needed to replicate the experimental observations. In that regard, it suffers in appearance from not seeming clinically relevant. For instance, we assume steady and laminar flow, whereas it is not the case *in vivo*. Recent work from INSIST partners^59^ identified up to 4 different patterns of flow in clotted regions of ischemic stroke patients: *slow anterograde, fast anterograde, retrograde* and mixed *anterograde-retrograde* flows. Anterograde flow designates the expected direction of the flow, *i*.*e*. from proximal to distal, while retrograde means the opposite direction. These different flow patterns undoubtedly affect the transport of lytic drugs to the occlusion site and, eventually, the evolution of the lysis. Also, the artery is here modeled by a rigid tube, and clot fragments stay in place until totally lysed, even when disconnected from the wall or the main clot chunk. Fortunately, thanks to its design, the model could be easily extended to include conditions closer to *in vivo* thrombolysis. We could instead reconstruct^60^ patient-specific geometries with collateral vessels and try to reproduce the complex flow conditions observed in AIS patients^59^. In doing so, we could hypothesize the effect of artery geometry, thrombus location, and drug injection patterns on lysis efficacy. Seners *et al*. have reported that distal thrombi lyse faster than proximal thrombi^61^. One could use the numerical model to assess whether this discrepancy comes from the shape and composition of distal clots, local flow conditions, or both. The model may estimate the risk of thromboembolism by accounting for the interaction of the disconnected clot components with the fluid. Using methods for moving boundaries or modifications of the PBB approach^62,63^, would allow to make these components move with the fluid, aggregate with other fragments, re-adhere to the main clot, or eventually leave the system, hence mimicking thromboemboli. Future works will incorporate these processes to extend the model to assess whether certain conditions in our numerical experiments are more likely than others to form emboli.

During the last two decades, we have witnessed an unprecedented revolution in treating acute ischemic stroke. Indeed, intravenous thrombolysis and endovascular mechanical thrombectomy therapies have entirely changed patient outcomes. However, disappointingly, while the endovascular mechanical thrombectomy allows for up to 85% of reperfusion in acute ischemia related to large vessel occlusion, the intravenous thrombolysis is still associated with low recanalization rates, less than 30%, in large and medium-sized vessels. Understanding conditions that increase lysis rates is essential to improve stroke therapies and should help explain clinical failures. In summary, we proposed a versatile and evolutive framework combining modeling and *in vitro* experiments to improve our understanding of fibrinolysis of pure fibrin occlusive clots in a tube. The one-dimensional model is purposely simplified to be solved analytically to predict front lysis time evolution. In future work, we aim to extend the model to incorporate more details of the clot structure and investigate further lysis patterns, including fragmentation, of clots with heterogeneous permeability and compositions. We intend to provide valuable knowledge to understand better the thrombus’s pathophysiology and its treatment with intravenous thrombolysis to improve the patient’s care.

## Data availability

The datasets used and/or analysed during the current study are available from the corresponding author on reasonable request.

## Acknowledgements

This research is part of the INSIST project^2^, which has received funding from the European Union’s Horizon 2020 research and innovation program under grant agreement No. 777072. This work was also supported by grants from the CHU Charleroi and the Fonds de la Chirurgie Cardiaque. The authors thank the contribution of Anaelle Taquin in the preparation and assistance during the experiments.

## INSIST Investigators

Charles Majoie^6^, Ed van Bavel^7^, Henk Marquering^6,7^, Nerea Arrarte-Terreros^6,7^, Praneeta Konduri^6,7^, Sissy Georgakopoulou^7^, Yvo Roos^8^, Alfons Hoekstra^9^, Raymond Padmos^9^, Victor Azizi^9^, Claire Miller^9^, Max van der Kolk^9^, Aad van der Lugt^10^, Diederik W.J. Dippel^11^, Hester L. Lingsma^12^, Nikki Boodt^10,11,12^, Noor Samuels^10,11,12^, Stephen Payne^13^, Tamas Jozsa^13^, Wahbi K. El-Bouri^13^, Michael Gilvarry^14^, Ray McCarthy^14^, Sharon Duffy^14^, Anushree Dwivedi^14^, Behrooz Fereidoonnezhad^15^, Kevin Moerman^15^, Patrick Mc Garry^15^, Senna Staessens^16^, Simon de Meyer^16^, Sarah Vandelanotte^16^, Francesco Migliavacca^17^, Gabriele Dubini^17^, Giulia Luraghi^17^, Jose Felix Rodriguez Matas^17^, Sara Bridio^17^, Bastien Chopard^1,2^, Franck Raynaud^1,2^, Rémy Petkantchin^1,2^, Vanessa Blanc-Guillemaud^18^, Mikhail Panteleev^19,20^, Alexey Shibeko^19^, Karim Zouaoui Boudjeltia^3. 6^Department of Radiology and Nuclear Medicine, Amsterdam UMC, location AMC, Amsterdam, the Netherlands; ^7^Biomedical Engineering and Physics, Amsterdam UMC, location AMC, Amsterdam, the Netherlands; ^8^Department of Neurology, Amsterdam UMC, location AMC, Amsterdam, the Netherlands; ^9^Computational Science Lab, Faculty of Science, Institute for Informatics, University of Amsterdam, Amsterdam, the Netherlands; ^10^Department of Radiology and Nuclear Medicine, Erasmus MC University Medical Center, PO Box 2040, 3000 CA Rotterdam, the Netherlands; ^11^Department of Neurology, Erasmus MC University Medical Center, PO Box 2040, 3000 CA Rotterdam, the Netherlands; ^12^Department of Public Health, Erasmus MC University Medical Center, PO Box 2040, 3000 CA Rotterdam, the Netherlands; ^13^Institute of Biomedical Engineering, Department of Engineering Science, University of Oxford, Parks Road, Oxford OX1 3PJ, UK; ^14^Cerenovus, Galway Neuro Technology Center, Galway, Ireland; ^15^College of Engineering and Informatics, National University of Ireland Galway, Ireland; National Center for Biomedical Engineering Science, National University of Ireland Galway, Ireland; ^16^Laboratory for Thrombosis Research, KU Leuven Campus Kulak Kortrijk, Kortrijk, Belgium; ^17^Laboratory of Biological Structure Mechanics, Department of Chemistry, Materials and Chemical Engineering “Giulio Natta”, Politecnico di Milano, Piazza Leonardo da Vinci 32, 20133 Milano, Italy; ^18^Institut de Recherches Internationales Servier, Coubevoie Cedex, France; ^19^Center for Theoretical Problems of Physicochemical Pharmacology RAS, Moscow, Russia; ^20^Dmitry Rogachev National Research Center of Pediatric Hematology, Oncology and Immunology, Moscow, Russia; Faculty of Physics, Lomonosov Moscow State University, Moscow, Russia.

## Author contributions statement

R.P., F.R. and B.C. designed and developed the models. R.P. wrote the article and the computer code for the models. A.R., K.Z.B., F.R. and B.C. designed the experiments, and A.R. and K.Z.B. performed them. All authors reviewed the manuscript.

