## Supplementary information for "A simplified mesoscale 3D model for characterizing fibrinolysis under flow conditions"

### A Permeability measurement of fibrin clots

We measured the outflow for two concentrations of fibrinogen, $2.4\,\mathrm{mg{\cdot}ml^{-1}}$ (Figure S1A) and $3.2\,\mathrm{mg{\cdot}ml^{-1}}$ (Figure S1B). Clots were formed in 800 $\mu$l total volume with $38.4-51.2$ $\mu$l of fibrinogen at $50\,\mathrm{mg{\cdot}ml^{-1}}$, Owren. 160 $\mu$l of thrombin was added to trigger clot formation during 15 minutes at 37°$C$. Clots have typical length of 2.6 cm, diameter 5.8 mm and final concentrations of fibrinogen of $2.4\,\mathrm{mg{\cdot}ml^{-1}}$ and $3.2\,\mathrm{mg{\cdot}ml^{-1}}$. Outflow was measured every ten minutes. As expected, outflows were constant over time, allowing us to measure clot permeability reliably. However, our results show high variability within the same experimental conditions (Figure S1C), highlighting how internal clot properties, in particular fiber radius, impact the permeability and, ultimately, the transport of proteins through the clot. Using Darcy's law, we estimated the permeability from the measurement of the outflow:

$$k = \frac{Q\mu}{\mathscr{A}} \frac{l_0}{\rho g (L - l_0)} \tag{S1}$$

where $\mathscr{A}$ is the section of the tube, $k$ the permeability, $\mu$ the dynamic viscosity, $L$ the length of the tube and $l_0$ the length of the clot.

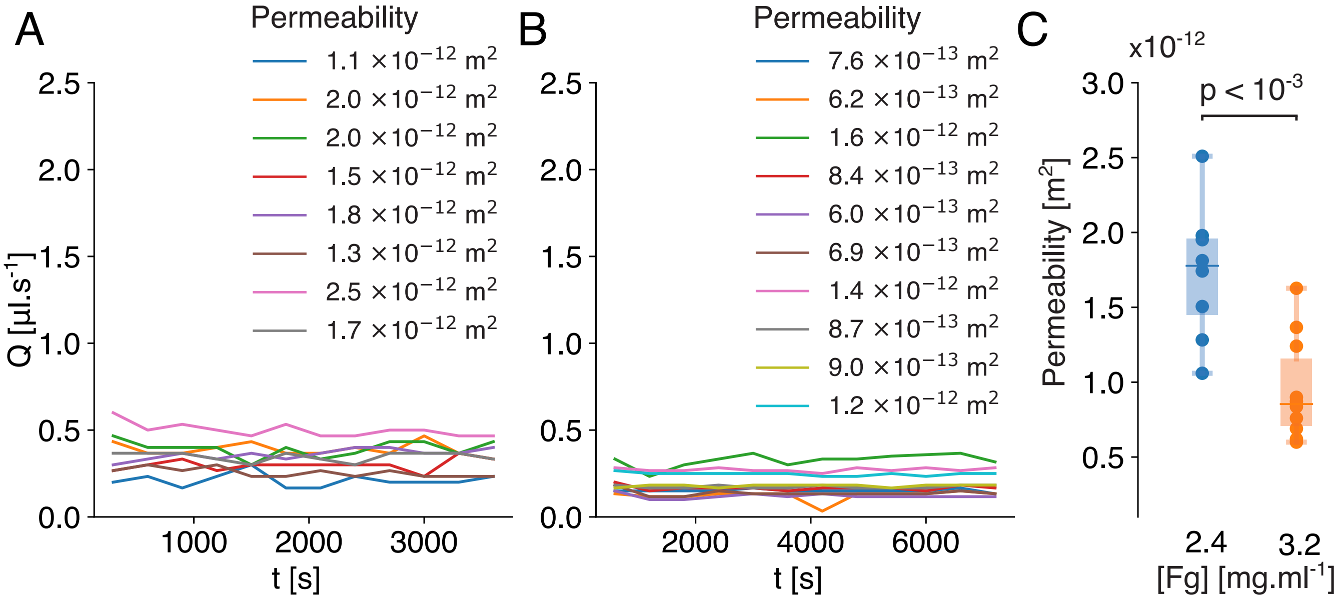


**Figure S1.** Permeability decreases with fibrinogen concentration. Outflow as function of time for fibrinogen concentration of **A**: $2.4\,\mathrm{mg{\cdot}ml^{-1}}$ and **B**: $3.2\,\mathrm{mg{\cdot}ml^{-1}}$. Each curve corresponds to a separate experiment, with the corresponding permeability indicated in the legend. **C**: Permeability as function of the concentration of fibrinogen. The boxplots represent the median as well as the first and third quartiles.

### B Lysis of the $n$-th slice with the analytical model

Before initiating the lysis of the second slice $x_1^\Delta$, the quantity $\overline{F}_0 \frac{u_f}{\Delta} t$ has to be advected over a distance $\Delta$. Hence, the anti-FA that accumulates in $x_1^\Delta$ is given by:

$$\overline{F}(x_1^\Delta, t) = \overline{F}_0 \frac{u_f}{\Delta} \left( t - \frac{\Delta}{u_f} \right) \cdot \Theta \left( t - \left( t_0 + \frac{\Delta}{u_f} \right) \right). \tag{S2}$$

The Heaviside step function $\Theta$ reflects that anti-FA does not reach $x_1^\Delta$ until $x_0^\Delta$ has been lysed, and the anti-FA has been transported over the distance $\Delta$. Thus, at the time $t = t_0 + \frac{\Delta}{u_f}$, the concentration of anti-FA at $x_1^\Delta$ is $\overline{F}_0 \frac{u_f t_0}{\Delta}$.

After injecting (S2) into (1) and integrating, we have :

$$
\begin{aligned}
\int_{F_0}^{F^*} \frac{dF(x_1^\Delta,t)}{F(x_1^\Delta,t)} &= -k_1\overline{F}_0 \frac{u_\mathrm{f}}{\Delta} \int_0^{t_1} \left(t - \frac{\Delta}{u_\mathrm{f}}\right) \Theta\left(t - \left(t_0 + \frac{\Delta}{u_\mathrm{f}}\right)\right) dt \\
&= -k_1\overline{F}_0 \frac{u_\mathrm{f}}{\Delta} \int_{t_0+\frac{\Delta}{u_\mathrm{f}}}^{t_1} \left(t - \frac{\Delta}{u_\mathrm{f}}\right) dt \\
\Rightarrow \log\left(\frac{F^*}{F_0}\right) &= -k_1\overline{F}_0 \frac{u_\mathrm{f}}{2\Delta} \left[t^2 - \frac{2\Delta}{u_\mathrm{f}} t\right]_{t_0+\frac{\Delta}{u_\mathrm{f}}}^{t_1} \\
&= -k_1\overline{F}_0 \frac{u_\mathrm{f}}{2\Delta} \left(t_1^2 - \frac{2\Delta}{u_\mathrm{f}} t_1 - \left(t_0 + \frac{\Delta}{u_\mathrm{f}}\right)^2 + \frac{2\Delta}{u_\mathrm{f}}\left(t_0 + \frac{\Delta}{u_\mathrm{f}}\right)\right),
\end{aligned}
$$

which after rearrangement gives:

$$
\left(t_1 - \frac{\Delta}{u_\mathrm{f}}\right)^2 - t_0^2 + \frac{2\Delta \log\left(\frac{F^*}{F_0}\right)}{k_1\overline{F}_0 u_\mathrm{f}} = 0. \tag{S3}
$$

Keeping the positive solution of the resulting quadratic equation, we get the time $t_1$ at which the second slice $x_1^\Delta$ is lysed:

$$
t_1 = \frac{\Delta}{u_\mathrm{f}} + \sqrt{t_0^2 + \frac{2\Delta}{k_1 u_\mathrm{f} \overline{F}_0} \log\left(\frac{F_0}{F^*}\right)}. \tag{S4}
$$

By applying the same reasoning recursively, we get the formula for the time to lyse $n$ slices :

$$
t_n = \frac{\Delta}{u_\mathrm{f}} + \sqrt{t_{n-1}^2 + \frac{2\Delta}{k_1 u_\mathrm{f} \overline{F}_0} \log\left(\frac{F_0}{F^*}\right)}.
$$

### C Analytical model independent of $\Delta$

We show in Figure S2 that the analytical formula for the lysis time of the thrombus does not depend on the choice of the spatial slicing $\Delta$, provided that $\Delta \ll L$.

We demonstrate it here analytically, starting with equation (5) and rearranging terms:

$$
\begin{aligned}
t_n &= \frac{\Delta}{u_\mathrm{f}} + \sqrt{t_{n-1}^2 + \frac{2\Delta}{k_1 u_\mathrm{f} \overline{F}_0} \log\left(\frac{F_0}{F^*}\right)} \\
&= \frac{\Delta}{u_\mathrm{f}} + \sqrt{t_{n-1}^2 + \beta\Delta} \\
\Rightarrow \left(t_n - \frac{\Delta}{u_\mathrm{f}}\right)^2 &= t_{n-1}^2 + \beta\Delta \\
\Rightarrow t_n^2 - 2\frac{\Delta}{u_\mathrm{f}} t_n + \frac{\Delta^2}{u_\mathrm{f}^2} &= t_{n-1}^2 + \beta\Delta.
\end{aligned}
$$

By taking the first order approximation of $\Delta$, we neglect the second order term $\frac{\Delta^2}{u_\mathrm{f}^2}$, which yields:

$$
t_n^2 - t_{n-1}^2 = 2\frac{\Delta}{u_\mathrm{f}} t_n + \beta\Delta. \tag{S5}
$$

On the other hand, we can write :

$$
t_n^2 - t_{n-1}^2 = (t_n - t_{n-1})(t_n + t_{n-1}) \approx 2t\,dt, \tag{S6}
$$

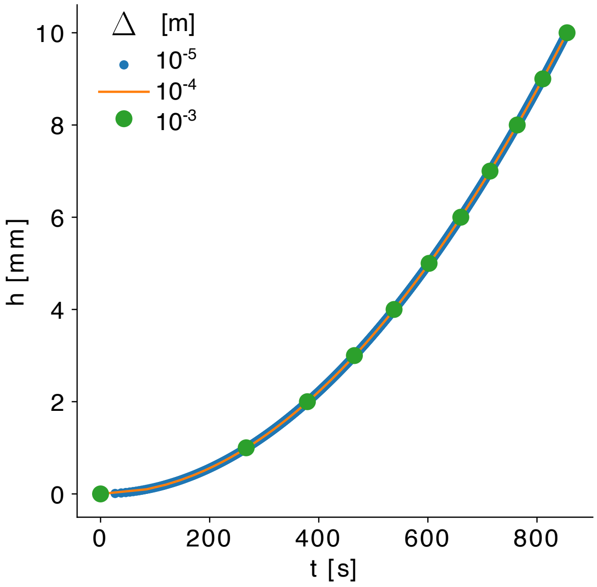


**Figure S2.** Lysis profiles of a 1 cm thrombus predicted with the analytical formula. $F_0 = 3.2$ mg·ml$^{-1}$, $\overline{F}_0 = 10$ ng·ml$^{-1}$, $\nabla P = 10^6$ Pa·m$^{-1}$, $k_1 = 12.5(\text{mg·ml}^{-1})^{-1}$. The lysis time predicted with the analytical formula is independent of the slicing of the thrombus Δ, is Δ is small compared to $L$.

where $dt$ is the variation of lysis time at time $t$ for the slice Δ. At first order in Δ, $t_n + t_{n-1} = 2t$ is a sufficient approximation. Combining (S5) and (S6), and writing $\Delta = dh$ we get:

$$2t\,dt = \left(\frac{2}{u_\mathrm{f}}t + \beta\right) dh \tag{S7}$$

$$\Rightarrow \frac{2t}{\frac{2}{u_\mathrm{f}}t + \beta} dt = dh \tag{S8}$$

$$\Rightarrow \left(u_\mathrm{f} - \frac{\beta}{\frac{2}{u_\mathrm{f}^2}t + \frac{\beta}{u_\mathrm{f}}}\right) dt = dh. \tag{S9}$$

By integrating, and setting $h = 0$ at $t = 0$ we finally get:

$$h(t) = u_\mathrm{f}t - \frac{1}{2}\beta u_\mathrm{f}^2 \log\left(2t/u_\mathrm{f}^2 + \beta/u_\mathrm{f}\right) + \frac{1}{2}\beta u_\mathrm{f}^2 \log\left(\beta/u_\mathrm{f}\right), \tag{S10}$$

which, using the properties of the log, can be rewritten as:

$$h(t) = u_\mathrm{f}\left(t - \frac{1}{2}\beta u_\mathrm{f} \log\left(\frac{2}{\beta u_\mathrm{f}}t + 1\right)\right), \tag{S11}$$

or:

$$h(t) = u_\mathrm{f}t - \frac{u_\mathrm{f}\log\left(\frac{F_0}{F^*}\right)}{k_1\overline{F}_0} \log\left(\frac{k_1\overline{F}_0}{\log\left(\frac{F_0}{F^*}\right)}t + 1\right), \tag{S12}$$

and which is independent of the slicing.

### D Experimental lysis fronts with various concentrations.

We put at disposition here lysis front measurements for fibrinolysis experiments that we conducted, while varying initial fibrin concentration (Figure S3A), and the tPA solution concentration (Figure S3B). The dots represent the experimental

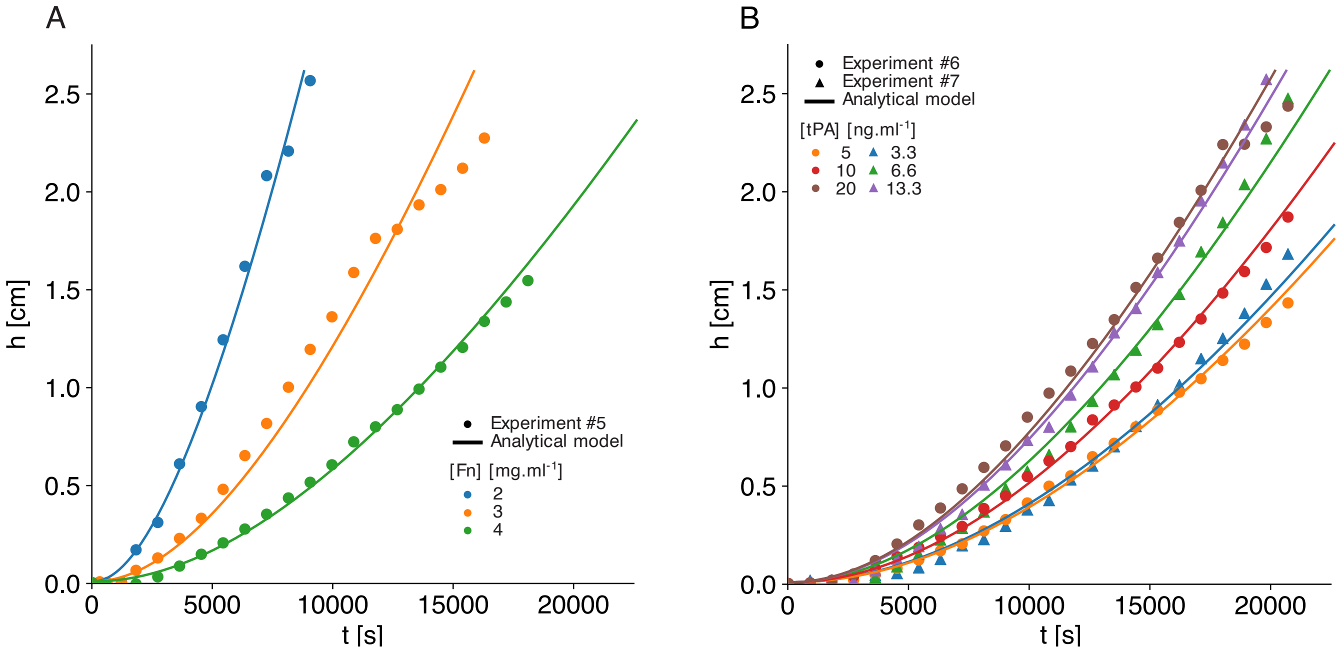


**Figure S3.** **A**: Comparison of lysis fronts obtained with the second set of experiments where fibrin concentration is varied, and the analytical model uses optimized reaction parameter obtained by minimizing the RMSE. tPA concentration is $10\,\mathrm{ng{\cdot}ml^{-1}}$. **B**: Comparison of lysis fronts obtained with the second set of experiments where tPA concentration is varied, and the analytical model uses optimized reaction parameter. We differentiate here two batches of experiments (dots, triangles) that might have suffered slightly different environmental conditions. Fibrin concentration is $3.2\,\mathrm{mg{\cdot}ml^{-1}}$.

measurements made at 15 minutes interval, and the solid lines the analytical model, assuming constant flow throughout the lysis, estimated with Darcy's law and Davies' equation for the thrombi permeability.

### E Pressure boundary conditions

Pressure boundary conditions (BCs) were imposed on both ends of the tube by imposing densities, as is usually done in LB. The 3D Zou-He pressure BC[1] and a novel pressure BC were tested. The novel BC showed better simulation stability when increasing the pressure difference; that is why it was chosen for the numerical model. This novel BC can also simulate a pulsatile pressure profile by applying a time modulation to the inlet density. Figure S4 presents an explanation of how the populations at the boundaries are computed.

Let us show how the density is imposed with this pressure BC. We consider the situation displayed in Figure S4, where we need to impose the density on the outlet boundary node $N$. After having copied the unknown populations $f_i$ from node $N-1$, the density at node $N$ can be computed as usually in LB:

$$\rho_{\mathrm{tmp}} = \sum_j f_j(\mathbf{x_N},t) + \sum_i f_i(\mathbf{x_N},t) = \sum_k f_k(\mathbf{x_N},t) \tag{S13}$$

where the $f_j$ designates the known populations from the streaming step, at node $N$. The equilibrium function $f_k^{eq}$ at node $N$, used in the standard BGK collision, reads[2]:

$$f_k^{eq}(\mathbf{x_N},t) = w_k\,\rho_{\mathrm{tmp}}\left(1 + \frac{\mathbf{u_N}\cdot\mathbf{c_k}}{c_s^2} + \frac{(\mathbf{u_N}\cdot\mathbf{c_k})^2}{2c_s^4} - \frac{\mathbf{u_N}\cdot\mathbf{u_N}}{2c_s^2}\right) \tag{S14}$$

where $w_k$ are the weighting factors, $\mathbf{c_k}$ the lattice discrete velocities, and $c_s$ the speed of sound magnitude in the lattice. By using the property that $\sum_k f_k^{eq}(\mathbf{x_N},t) = \sum_k f_k(\mathbf{x_N},t) = \rho_{\mathrm{tmp}}$ at node $N$, we subtract and add the following equilibrium functions to the populations $f_k$ at node $N$:

$$\begin{aligned}\rho' =&\sum_k f_k(\mathbf{x_N},t) - f_k^{eq}(\rho_{\text{tmp}},\mathbf{u_N})\\&+ f_k^{eq}(\rho_{\text{target}},\mathbf{u_N})\\=&\sum_k f_k(\mathbf{x_N},t) - \sum_l f_l^{eq}(\rho_{\text{tmp}},\mathbf{u_N})\\&+\sum_m f_m^{eq}(\rho_{\text{target}},\mathbf{u_N})\\=&\rho_{\text{tmp}} - \rho_{\text{tmp}} + \rho_{\text{target}}\\=&\rho_{\text{target}}\end{aligned}$$

By juggling with these equilibrium functions, we can thus impose the density $\rho_{\text{target}}$ at the boundary. It is worth noting that the velocities $\mathbf{u_N}$ remain unchanged.

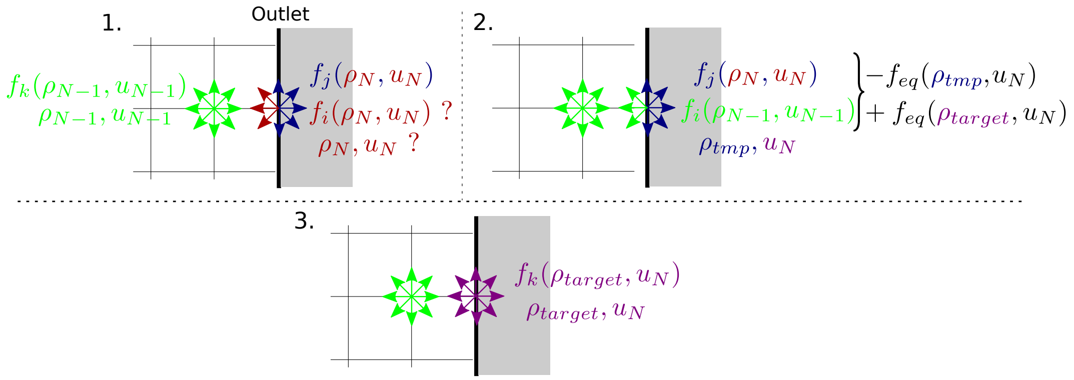


**Figure S4.** Pressure boundary condition used for inlet and outlet, with $\rho_N$ the density and $u_N$ the velocity at node $N$. The computation steps are the following (here showing outlet): **1**. Pre-computation state: At node $N$, populations $f_j$ are known from the streaming of neighbor nodes at previous iterations, and populations $f_i$, density, and velocities are unknown. At node $N-1$, $f_k$, density and velocities are known. It is chosen as the previous node in the direction normal to the boundary. **2**. Unknown populations at node $N$ are copied from populations at node $N-1$. From populations $f_j$ and $f_i$, a temporary density $\rho_{\text{tmp}}$ and the velocities are computed. Two equilibrium functions are then computed, one with $\rho_{\text{tmp}}$, and the other with the target density $\rho_{\text{target}}$. We subtract the former and add the latter to the populations at node $N$. **3**. Node $N$ now has target density $\rho_{\text{target}}$ and populations $f_k$.

### F Particles transport

The particles in the numerical model embody the *anti-FA* entity, which accounts for the lysis of fibrin. They are subject to *advection*, the transport driven by the fluid (lytic solution), and *diffusion*, which is the movement due to thermal agitation and molecular collisions.

#### F.1 Advection

The Palabos library offers a built-in algorithm that automatically transports particles advected by the fluid. However, it is a one-size-fits-all solution and is not necessarily the most efficient way to advect the particle for a specific model. Looping over all particles at each iteration step is costly. In our thrombolysis model, the flow varies very slowly during most of the lysis. It is thus interesting in our case, to increase the advection time scale according to the flow velocity. Practically, instead of moving on a distance $dx$ the particles at each lattice-Boltzmann iteration, we move them of $T dx$ after $T$ iterations, where $T$ is computed such that the particles move at most of half a lattice site:

$$T = \frac{0.5}{\text{maximum velocity inside domain}(t)}. \tag{S15}$$

This computational trick allows a speedup of around a factor 10 in a typical lysis simulation, without any visible difference on the lysis curves. The advection velocity of each particle is computed as with the Palabos solver, using weighted averaging of the velocity at the 4 closest lattice nodes.

#### F.2 Diffusion

The Péclet number:

$$\mathrm{Pe} = \frac{uL}{D} \tag{S16}$$

where $u$ is the mean velocity of the fluid, $L$ is a characteristic dimension, and $D$ the diffusion coefficient of the diffusing mass in the medium, gives an approximation of the ratio of advection against diffusion. For high Péclet ($\mathrm{Pe} \gg 1$), diffusion can be neglected but should otherwise be taken into account. In the clotted tubes of the experiments presented in the manuscript, and for tPA molecules, the initial Péclet is:

$$\mathrm{Pe}_i = \frac{u_{\mathrm{solution}}\, d_{\mathrm{tube}}}{D_{\mathrm{tPA}}} \sim \frac{3 \cdot 10^{-7} \cdot 10^{-2}}{3.3 \cdot 10^{-11}} \sim 100 \tag{S17}$$

which shows that the diffusion of tPA is non-negligible to initiate the lysis of the clot.

Since the anti-FA is modeled by particles, the most straightforward way to implement their diffusion is by individually giving a random velocity at each simulation time step $dt$, of norm :

$$|v_{\mathrm{Diff}}| = \frac{\sqrt{2nD_{\mathrm{tPA}}dt}}{dt} \tag{S18}$$

where $n = 3$ is the spatial dimension of the model.

It can be verified that this method yields the expected collective behavior, by computing the mean squared displacement (MSD) of the $N$ particles:

$$\mathrm{MSD}(t) = \frac{1}{N}\sum_{i=1}^{N} |\mathbf{x_i}(t) - \mathbf{x_i}(0)|^2 \tag{S19}$$

where $\mathbf{x_i}(t)$ is the 3D position of particle $i$ at time $t$.

It can be shown that the MSD approximates the diffusion coefficient:

$$\mathrm{MSD}(t) = 2nD_{\mathrm{tPA}}t. \tag{S20}$$

By simulating $N = 100$ particles in a system with no advection, already good agreement can be observed with eq. (S20), as shown in Figure S5.

### G Porosity and permeability validation

Let us here validate our implementation of the PBB model for computing flow in simulated fibrin clots.

Let us recall Darcy's law for laminar flow[3] :

$$u_{\mathrm{f}} = -\frac{k}{\rho \nu}\frac{\Delta P}{L}. \tag{S21}$$

with $\nu$ the kinematic viscosity, $\rho$ the fluid density, $\Delta P$ the pressure drop and $k$ the *permeability*.

Walsh et al. show that the permeability $k_{\mathrm{PBB}}$ in $\mathrm{m}^2$ of PBB voxels is related to $\gamma$ and the simulation time step $\Delta t$ as follows:

$$k_{\mathrm{PBB}} = \frac{(1-\gamma)\nu\,\Delta t}{2\gamma}. \tag{S22}$$

Combining (14), (S21) and (S22), we can express Darcy's law in terms of $\gamma$:

$$u_{\mathrm{f}} = -\frac{\Delta t}{2\gamma}\frac{\Delta P}{L\rho}. \tag{S23}$$

We measured the values of the fluid velocity $u_{\mathrm{f}}$ in numerical simulations, and compared with the analytical solution of Darcy and Walsh equation (S23) in Figure S6A. Flow was simulated in a 3D rectangular duct filled with a porous medium with bounce-back fractions of $\gamma = 0.2, 0.4, 0.8$ respectively. The velocity profile was measured along the $y$-axis (transversal) in the center of the duct. In the analytical solution, we imposed no-slip conditions, i.e. $u_{\mathrm{f}} = 0$ at the walls.

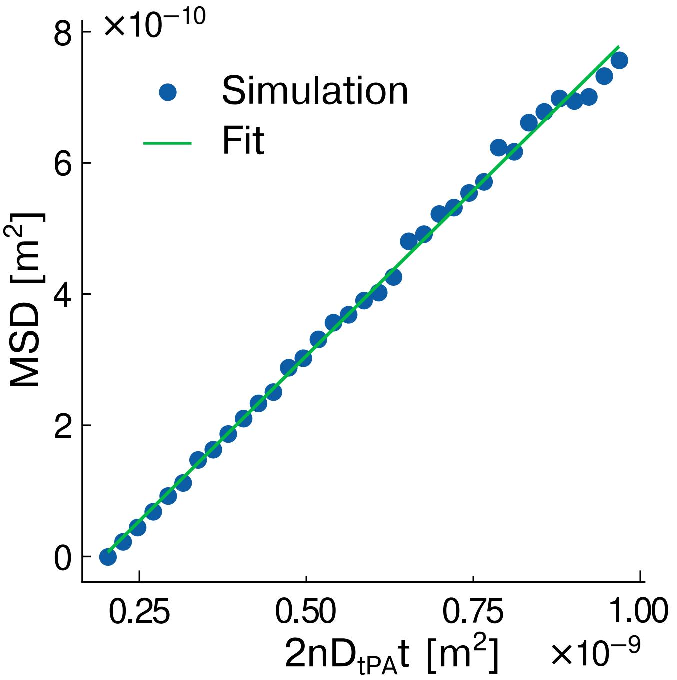


**Figure S5.** Diffusion of $N = 100$ anti-FA particles in a simulation without flow. A linear fit is shown between the MSD and the quantity $2nD_{\mathrm{tPA}}t$, which slope is 1.007, as expected from equation (S20).

Next, we investigate the validity of the PBB method for a typical range of fibrin clot permeabilities. Other commonly used laws to describe fibrous porous media permeabilities are Clague's equation[4], a numerical solution of permeability of randomly organized fibrous media:

$$k_{\mathrm{C}}(R_f, n_s^*) = 0.50941 R_f^2 \left( \left( \frac{\pi}{4n_s^*} \right)^{0.5} - 1 \right)^2 e^{-1.8042 n_s^*}, \tag{S24}$$

and Jackson-James' equation[5], the weighted average of the solution to the Stokes equation for flow parallel to or normal to 2D periodic square arrays of cylinders:

$$k_{\mathrm{JJ}}(R_f, n_s^*) = R_f^2 \frac{3}{20 n_s^*} (-\ln(n_s^*) - 0.931). \tag{S25}$$

Permeabilities measured with the model as a function of input solid fractions are shown in Figure S6B-D, and compared with their empirical counterparts equations (16),(S24) and (S25). The permeability values for the physiological range of low solid fractions ($0.003 - 0.04$, i.e. $1 - 4\,\mathrm{mg{\cdot}ml^{-1}}$ fibrinogen concentration) do not vary substantially from one model to another; thus, if the clots studied are highly porous, the choice of model has little impact. Clague's law predicts an increase of permeability with the increase of solid fraction in the $0.8 - 1$ range, and Jackson-James' law asymptotically approaches zero as $n_s^*$ approaches 0.4. Thus, in order to be able to simulate the whole range of solid fractions, Davies' permeability law was chosen for the model.

### H Deviation of numerical from analytical model

We show here the deviation of the mesoscopic numerical lysis front, from the front given by the analytical formula, in the conditions of Figure 4.The relative absolute difference $\varepsilon$ on the front position in time is shown on Figure S7. We see that the deviation stays below 5% during most of the lysis, and increases up to 20% in the last third of the total lysis time. This increase in the difference is due to the assumption of constant flow made in the analytical model.

### I Lysis of realistic heterogeneous clots

We generated numerical thrombi directly from ischemic stroke patient thrombi retrieved by thrombectomy and sliced by Staessens et al.[8]. Platelet stained slices are converted to grayscale using the 'L' algorithm of the Python Imaging Library.

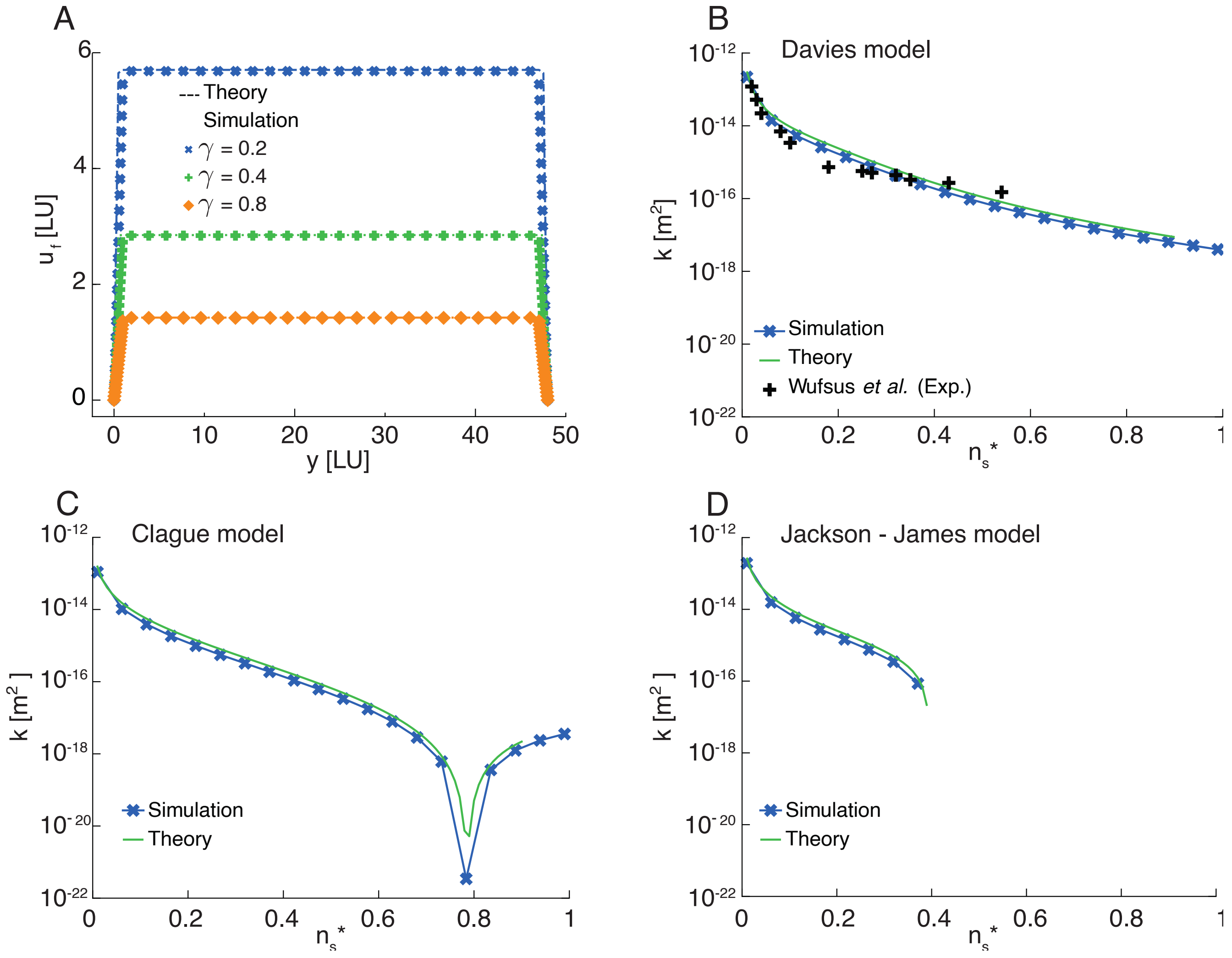


**Figure S6.** **A**: Comparison of fluid velocity of a transverse slice of a clot in a rectangular duct, between Darcy's law and our implementation of the PBB method, for three bounce-back fractions $\gamma = 0.2, 0.4, 0.8$. Velocity and position are given in lattice units (LU). Space and time steps were $\Delta x = 1.13 \cdot 10^{-4}$ m and $\Delta t = 1.83 \cdot 10^{-5}$ s, $\nu = 3 \cdot 10^{-6}$ m$^2$·s$^{-1}$, and the pressure gradient $\nabla P = 7700$ Pa·m$^{-1}$. The domain size was $N_x = N_y = N_z = 48$ lattice nodes. **B,C,D**: Fibrous thrombus intrinsic permeability $k$ as a function of fiber solid fraction $n_s^*$. Wufsus et al.[6] *in vitro* fibrin gels measurements (black +), Davies' equation (B)[7], Clague (C)[4] and Jackson and James (D)[5], according to equations (16),(S24) and (S25) respectively. Simulations were made in a tube of radius 4.5 mm (23 nodes), length 1 cm (46 nodes), with a fully occluding homogeneous thrombus (i.e., same dimensions as the tube). The space-time discretization was $\Delta x = 2.17 \cdot 10^{-4}$ m and $\Delta t = 1.58 \cdot 10^{-3}$ s, and the pressure gradient $\nabla P = 100$ Pa·m$^{-1}$ The hydrated fiber radius ranges from $R_f = 51$ to 60 nm for formulae and simulations, as in[6].

We then assign a fibrin concentration in a linear scale ranging from 1 mg·ml$^{-1}$ (white, no platelets) to 8 mg·ml$^{-1}$ (black, platelet-rich), Figure S8A. Platelet-rich zones are supposed to be more tPA-resistant and to have lower permeability[9],[10],[11], which we translate in the present model with higher fibrin concentration regions. We lysed a RBC-rich clot and a PLT-rich clot with the numerical model, as well as homogeneous clots with the same average concentration of 2.45 and 3.92 mg·ml$^{-1}$ respectively, Figure S8B. The lysis of the heterogeneous clots was slightly faster, and the recanalization occurred slightly before, Figure S8C.

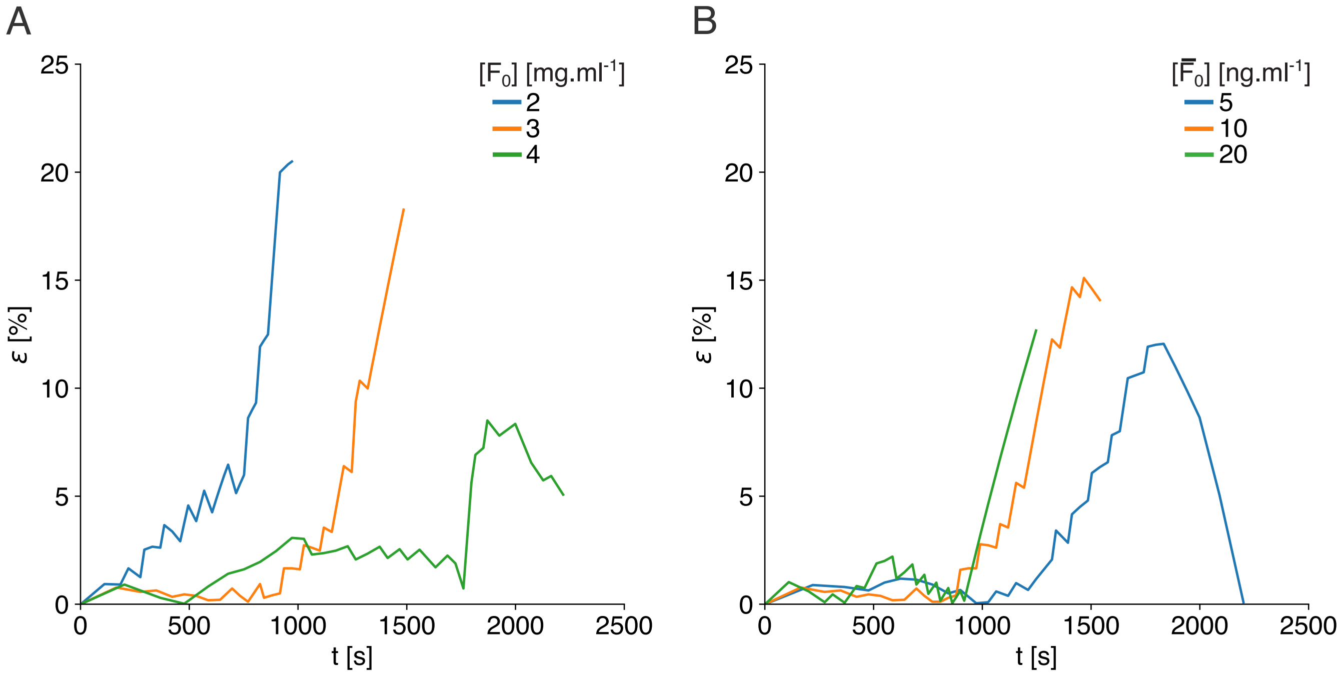


**Figure S7.** Lysis front position deviation in time, between the numerical 2D and analytical models, for different concentrations of fibrin (**A**) and anti-FA (**B**). Baseline parameters are $F_0 = 3\,\mathrm{mg{\cdot}ml^{-1}}$, $\overline{F}_0 = 10\,\mathrm{ng{\cdot}ml^{-1}}$, $k_1 = 280\,(\mathrm{s{\cdot}mg{\cdot}ml^{-1}})^{-1}$, $\nabla P_0 = 8250$ $\mathrm{Pa{\cdot}m^{-1}}$, $\Delta x = 1.14 \cdot 10^{-4}$ m, $\Delta t = 1.83 \cdot 10^{-4}$ s.

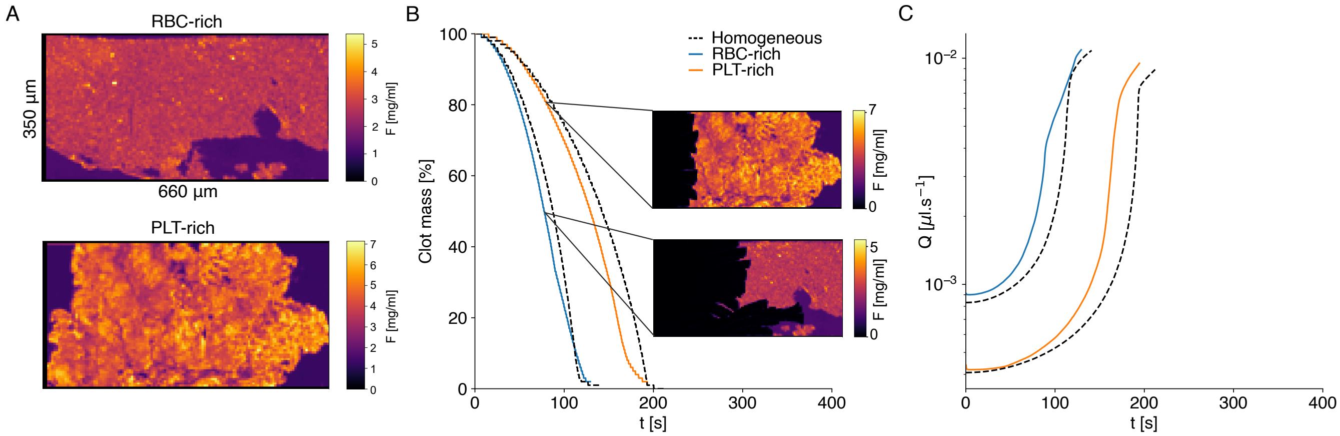


**Figure S8.** **A**: Platelet stained patient clot slices from[8] are digitalized by assigning a fibrin concentration in a linear scale, based on platelet staining. **B**: These clots are lysed with the numerical model, with an imposed pressure gradient $\nabla P = 8000\,\mathrm{Pa{\cdot}m^{-1}}$ and a reaction constant $k_1 = 1400\,(\mathrm{s{\cdot}mg{\cdot}ml^{-1}})^{-1}$, and compared against homogeneous clots with the same concentration. **C**: The mean throughput is monitored during the lysis. RBC-rich clots are more permeable since before the start of the lysis, the average fibrin concentration being lower than the PLT-rich's one.

3. Darcy, H. *Les fontaines publiques de la ville de Dijon* (Librairie des corps impériaux des ponts et chaussées et des mines,

1856), victor dalmont edn.
